# A Conserved TCRβ signature dominates a highly polyclonal T-cell expansion during the acute phase of a murine malaria infection

**DOI:** 10.1101/2020.07.27.222570

**Authors:** Natasha L. Smith, Wiebke Nahrendorf, Catherine Sutherland, Jason P. Mooney, Joanne Thompson, Philip J. Spence, Graeme J. M. Cowan

## Abstract

CD4_+_ αβ T-cells are key mediators of the immune response to a first *Plasmodium* infection, undergoing extensive activation and splenic expansion during the acute phase of an infection. However, the clonality and clonal composition of this expansion has not previously been described. Using a comparative infection model, we sequenced the splenic CD4_+_ T-cell receptor repertoires generated over the time-course of a *Plasmodium chabaudi* infection. We show through repeat replicate experiments, single-cell RNA-seq, and analyses of independent RNA-seq data, that following a first infection – within a highly polyclonal expansion – T-effector repertoires are consistently dominated by TRBV3 gene usage. Clustering by sequence similarity, we find the same dominant clonal signature is expanded across replicates in the acute phase of an infection, revealing a conserved pathogen-specific T-cell response that is consistently a hallmark of a first infection, but not expanded upon re-challenge. Determining the host or parasite factors driving this conserved response may uncover novel immune targets for malaria therapeutic purposes.

## 1. Introduction

Although protective natural immunity against clinical malaria is slow to develop and requires years of repeated exposure (1), protection against severe disease is obtained after a more limited number of symptomatic infections (2,3). The acquisition of this naturally acquired immunity is mediated by both antibody (reviewed in (4,5)) and T-cell responses (6); the latter being crucial for B-cell class switching and affinity maturation. As well as guiding the humoral response, CD4_+_ T-cells play a key role in restricting the growth and pathogenesis of blood-stage *Plasmodium* through cytokine secretion and macrophage activation (reviewed in (7)). However, the antigenic drivers and developmental dynamics underlying this naturally acquired immunity remain poorly understood, presenting major challenges for effective vaccine design.

In animal models of malaria, a *Plasmodium* infection in previously unexposed individuals initially produces a massive expansion of CD4_+_ T-cells in the spleen (8,9), a major site of the developing immune response (10). The size of this response, together with the generation of a highly diverse range of cellular responses, suggests that the splenic expansion of CD4_+_ populations is highly polyclonal, as opposed to the expansion of a minor (oligoclonal) subset of the repertoire. However, it is not known whether this expansion is primarily a non-specific response, such as a result of cytokine-driven bystander activation, or whether it is dominated by antigen-specific responses generated through classic TCR-engagement mediated clonal expansion. Spectratyping (CDR3 length analysis) of T-cell receptor (TCR) β chain repertoires induced by the rodent malaria *Plasmodium berghei* has previously detected a unique TCRβ CDR3 length signature enhanced over the course of infection, suggesting that there may be a clonal response to specific antigenic peptides (11). In agreement with this, an expanded fraction of CD4_+_ T-cells and fast-responding cytokine secretors that respond to a secondary challenge has been observed following a *Plasmodium chabaudi (AJ)* infection in mice, indicating initial priming by the parasite, and the presence of pathogen-specific T-cells within the CD4_+_ T-cell population (9). Alternatively, there is evidence from *P. falciparum*, that the PfEMP1 binding domain, CDR1a, stimulates CD4_+_ T-cells non-specifically through TCR-independent pathways (12), and that regulatory T-cell (Treg) proliferation during an infection can be induced in an antigen non-specific manner (13). Non-specific proliferation of T-cells due to cross-reactivity in response to *P. falciparum* antigens has also been reported (14).

Overall, proliferation is likely to be a combination of activation dynamics. However, whether a detectable clonal malaria-specific CD4_+_ T cell response that is conserved between individuals, and thus a potential focal target for therapeutics, is induced, has not previously been demonstrated.

Advances in high-throughput TCR repertoire sequencing techniques now allow deep profiling of immune responses. This approach has been used to ascertain clonality of T-cell responses, identify expanded T-cell clones and determine if conserved or ‘public’ responses between individuals are generated following antigenic stimulation (reviewed in (15)). Repertoire sequencing thus provides a novel, immune-focused approach to delineate the clonality of the developing immune response to a malaria infection.

Here, using bulk TCRβ repertoire sequencing we examine the dynamics and clonal structure of the splenic CD4_+_ T-cell repertoires generated during infection with the well-established mouse malaria model *Plasmodium chabaudi (AS)*. By comparing serially blood passaged (SBP) and recently mosquito-transmitted (MT) *P. chabaudi* infections, Spence *et al*. (2013) demonstrated that vector transmission of *P. chabaudi* intrinsically modified parasite gene expression in asexual blood-stage parasites, eliciting an altered host immune response that in turn regulates parasite virulence. In this model, infection with SBP parasites leads to hyperparasitaemia with more severe disease during the acute phase of infection. In contrast, mosquito transmission (MT) attenuates parasite growth and virulence, through a mechanism associated with epigenetic reprogramming of the expression of the subtelomeric multigene families, including the variant surface antigen (VSA) family. We have used this comparative model to compare TCR repertoires integral to an immune response that rapidly controls parasite growth, against a less effective response that fails to control parasite replication and induces immunopathology (8,16). We have sequenced the T-naive (T_N_), T-effector (T_E_), T-effector memory (T_EM_) and T-central memory (T_CM_) CD4_+_ splenic TCRβ repertoires elicited in mice over the time-course of both MT and SBP *P. chabaudi* infections. We report, for both infection types, that the T_E_ expansion seen during the acute phase of a *P. chabaudi* infection is highly polyclonal. However, within this diverse expansion, a conserved pathogen-specific response characterised by TRBV3 gene usage consistently dominates the effector repertoire following a first infection, and we further profile this response using single-cell RNA-seq.

## 2. Materials and Methods

### 2.1 Mice infections

All procedures were carried out in accordance with UK Home Office regulations (Animals Scientific Procedures Act, 1986; project licence number 70/8546 and P04ABDCAA) and were approved by the Ethical Review Body of the University of Edinburgh. C57Bl/6 mice were bred and housed under specific pathogen free conditions at the University of Edinburgh and subjected to regular pathogen monitoring by sentinel screening. Mice were housed with at least one companion in individually ventilated cages furnished with autoclaved woodchip, fun tunnel and tissue paper at 21 ± 2°C under a reverse light-dark cycle (light, 19.00 – 07.00; dark, 07.00 – 19.00) at a relative humidity of 55 ± 10%. *P. chabaudi (AS)* parasites were obtained from the European Malaria Reagent Repository at the University of Edinburgh. 8-10wk old C57Bl/6 female mice were infected with *P. chabaudi (AS)* by intra-peritoneal injection of 1×10_5_ parasitised erythrocytes that had either been maintained by serial blood-passage over a high number of generations (SBP) or undergone a single passage following mosquito transmission (MT) as per Spence *et al* (2012). Each transmission group consisted of 5 cages of 5 mice, with 5 unchallenged mice from the same cohort used as experimental controls. Mice (n=5 per transmission group) were euthanased on days 6, 10, 20, 40 and 60 post-infection. In a repeat experiment, mice (n=4 per time-point) were infected with MT *P. chabaudi (AS)-GFP* (17) by intra-peritoneal injection of 1×10_5_ parasitised erythrocytes, and were euthanased at days 4, 7, 11 and 14 post-infection. For the re-challenge experiment, mice were infected with MT *P. chabaudi (AS)-GFP* by intra-peritoneal injection of 1×10_5_ parasitised erythrocytes. Mice (n=4) underwent a homologous re-challenge at day 60 post infection and were euthanased 7 days post re-challenge (day 67). To eliminate chronic infection before re-challenge, 0.288mg/ml of chloroquine diphosphate salt (Sigma), supplemented with glucose for palatability, was dissolved in drinking water daily for 10 days, from day 30 to day 40 post-infection (18). For the single-cell RNA-seq experiment, mice (n=2) were infected with MT *P. chabaudi (AS)-GFP* by intra-peritoneal injection of 1×10_5_ parasitised erythrocytes, and euthanased at day 7 post-infection.

### 2.2 Cell sorting

CD4_+_ splenic T-cell populations of interest were isolated by FACS using a BD FACSAria III instrument, according to gates described by Spence *et al*. (2013): T_N_ (CD62L_+_ CD127_+_), T_E_ (CD62L_-_ CD127_-_), T_EM,_ (CD44HI CD127_+_ CD62L_-_), and T_CM_ (CD44HI CD127_+_ CD62L_+_). For the repeat and re-challenge experiments, only T_E_ and T_N_ populations were isolated. Cells were sorted into 50μL FACS buffer and stored at −80°C until processing. For the single-cell experiment, CD3_+_ CD4_+_ splenocytes were sorted in to 100μL 0.04% BSA in PBS, to generate the single-cell suspension required for 10x Genomics sequencing.

### 2.3 Unbiased bulk TCR amplification & sequencing

RNA was extracted from CD4_+_ splenic T-cell populations of interest using Dynabeads mRNA purification kit (ThermoFisher Scientific). cDNA was synthesised from each RNA preparation by adding the following to each sample: 4μL First-strand Buffer, 2μL 10mM dNTP mix, 2μL 20mM DTT and 2μL SMART-PTO2 oligo (5’ AAGCAGTGGTATCAACGGAGAGTACATrGrGrG_(3)_ 3’), 0.5μL RNase inhibitor (Clontech 2313A), and 2μL (100U/μL) SMARTScribe reverse transcriptase (Clontech). For the repeat experiments, unique molecular identifiers (UMIs) (19) were incorporated during cDNA synthesis by replacing the template-switch oligo with 2μL SMARTNNN oligo (5’AAGCAGUGGTAUCAACGCAGAGUNNNNUNNNNUNNNNUCTTrG_(3)_3’). Samples were then incubated at 42°C for 70 minutes, before the reaction was terminated by heating at 70°C for 10 minutes. cDNA synthesised with SMARTNNN oligos were treated with 1 μl of Uracil DNA glycosylase (5 U/μl, New England Biolabs) and incubated for 15 minutes at 37°C. PCR was then used to generate TCRβ V-region amplicons, using indexed forward primers composed of the SMART synthesis oligo sequence fused to a P7 Illumina tag, and a reverse primer within the TCR-C region fused to a P5 Illumina tag (P5-mTCRBrev3: 5’ AATGATACGGCGACCACCGAGATCTACACCTTGGGTGGAGTCACATTTCT 3’). Amplified products were purified by extraction from excised agarose gel bands. Single-end 1×400bp and asymmetric 100bp+400bp (to incorporate UMIs) sequencing was performed on an Illumina MiSeq platform.

### 2.4 Bulk TCR repertoire analyses

Bulk TCR sequence data was initially processed using MiGEC (20) and MiXCR (21) software with default settings. Samples were excluded from further analyses if the repertoire contained fewer than 10,000 total reads after processing, as this was indicative of poor sample preservation or preparation. A combination of custom pipelines of Python scripts and VDJtools software (22) was used to analyse and plot the MiXCR output. Statistical analyses were performed using SciPy Python software (23). A TCR clone was defined by 100% amino acid sequence identity of the CDR3 region, and IMGT nomenclature used for gene usage. Only in-frame (functional) CDR3s were analysed. A modified version of the Swarm algorithm (24) was used to cluster highly homologous CDR3 amino sequences, with identical V-gene usage, within 1 amino-acid mismatch of each other. The Gliph2 package (25) was also used to identify enriched amino acid motifs within the contact region of CDR3 sequences; the unchallenged T-naïve repertoires were used to make custom murine reference files for this. Network analyses was undertaken using Gephi (26) software (v0.9.2). Generation probability of TCRs (Pgen) was calculated using OLGA (27).

### 2.5 Publicly available RNA-seq data analyses

Raw FASTQ files from RNA-seq data obtained from the spleens of C57Bl/6 mice infected with blood-stage *Plasmodium chabaudi (AS)* and *Plasmodium chabaudi (CB)* were downloaded from the ArrayExpress archive: ENA – ERP004042 and ENA – ERP005730 respectively. For comparison of infection with other pathogens, raw FASTQ files of RNA-seq data obtained from whole blood of C57Bl/6 mice infected with a variety of pathogens, were downloaded from the NCBI short read archive, under accession SRR7821557. Normalised Trbv3 gene expression values were also obtained from Singhania *et al*. (2019). RNA-seq data was aligned using MiXCR (21) software, and a combination of custom pipelines of Python scripts and VDJtools software (22) was used to analyse and plot the MiXCR output.

### 2.6 Single-cell sequencing, data processing and analyses

Two barcoded cDNA libraries were prepared from sorted samples using the Chromium Single Cell 5’ Library Kit v3.1 (28). Full length V(D)J segments were enriched from amplified cDNA with primers specific to the TCR constant region using the Chromium Single Cell V(D)J Enrichment Kit – Mouse T-Cell. Sequencing was performed using the High-Output v2.5 Kit on a NextSeq 550 platform. Initial processing of sequence files, including mapping of reads to the mouse reference genome (GRCm38), generation of count matrixes and assembly of TCR alpha and beta chains, was carried out using CellRanger 3.1.0. To exclude potential multiplets, poor quality cells or non T-cells, single T-cells were identified by the expression of a single productive beta chain. Barcodes lacking a beta chain or assigned to multiple were excluded, leaving data from 3333 single T-cells (1658 and 1675 from mouse 1 and 2 respectively). Downstream analyses were performed in R using Seurat 3.1.5 (29). Genes expressed in fewer than 3 cells, as well as all *Trav/j* and *Trbv/d/j* genes were excluded. Cells expressing fewer than 200, or over 3000 genes and/or more than 5% mitochondrial genes were removed. The filtered matrix was normalised using Seurat’s LogNormalize with default parameters and the top 2,000 variable genes were identified using the FindVariableFeatures ‘vst’ method, before centring and scaling of the matrix. Dimensionality reduction by PCA was carried out and the top 30 principal components were used as input for graph-based clustering. Clusters were visualised by UMAP. A small, poorly *Cd4*-expressing cluster was identified, and these cells were excluded as contaminants. The above normalisation and clustering steps were repeated with the remaining 2976 cells (1491 and 1485 from mouse 1 and 2 respectively). Differential gene expression analysis using the Wilcoxon rank sum test through FindAllMarkers was used to identify marker genes for each cluster.

## 3. Results

Mice were infected with *P. chabaudi (AS)* parasitised erythrocytes from donor mice infected with either recently MT or SBP parasites, and followed for 60 days of infection. For repeat experiments, only MT parasites were used, as these represent a less artificial experimental model. At each time point, spleens were harvested and CD4_+_ splenic T-cells populations of interest were isolated by FACS before RNA was extracted, reverse transcribed, and TCRβ chains amplified and sequenced.

### 3.1 TRBV3 gene usage dominates a highly polyclonal T-effector expansion

Consistent with previously published data for *P. chabaudi* (8,30), CD4_+_ splenic T-effectors (T_E_) were maximally expanded during the acute phase of infection, increasing by up to 10-fold. Expansion coincided with the peak of parasitaemia before contracting back to pre-challenged levels between day 20 and 40 post-infection (Figure 1A, 1B). We first hypothesised that if the T_E_ expansion in the acute phase of infection was solely the result of non-specific activation, V gene usage and V/J allele usage within the T_E_ repertoire would mirror that of T_N_ repertoires, despite the vast cellular proliferation. Thus, there would be no change in the distribution of V or V/J allele usage post parasite-challenge. However, a distinct increase in TRBV3 gene usage observed during the acute phase of infection, differentiates challenged T_E_ repertoires from both unchallenged T_N_ and T_E_ repertoires, and from challenged T_E_ repertoires at later time-points (Figures 2A, 2B, Supplementary Figure 1). For MT infections during the acute phase, TRBV3 encodes on average 23.7% (± 2.03 95% CI) of the effector repertoire at day 6 and 21.6% (± 2.21 95% CI) at day 10 post-infection, compared to only 7.6% (± 0.47, 95% CI) of the unchallenged naïve repertoire. This finding was repeated in a second independent experiment (Supplementary Figure 2), and through analysis of publicly available RNA-seq data for *P. chabaudi (AS)* and *P. chabaudi (CB)* (accession E-ERAD-221 and E-ERAD-289 respectively, Supplementary Figure 3) when comparing unsorted TCRβ repertoires. This increase in TRBV3 usage was more delayed in SBP challenged repertoires, not apparent until day 10 post-challenge, where at its peak it encoded 17.2% (± 2.54, 95% CI) of the T_E_ repertoire.

An overall strong positive correlation between challenged T_E_ and unchallenged T_N_ V/J allele usage was evident during the effector expansion and indicates a highly polyclonal response with broad expansion of the naïve precursor pool. However, three specific TRBV3/J allelic combinations,TRBV3-TRBJ1-1, TRBV3-TRBJ2-4 and TRBV3-TRBJ2-7, deviated from this trend and are disproportionately increased in challenged T_E_ repertoires of mice infected with MT parasites at both days 6 and day 10 (Figure 2Ci), and for SBP infections by day 10 post-infection (Figure 2Cii). This specific V/J usage was conserved across all individual replicate mice infected with MT parasites during the acute phase of infection, though the response is more varied between SBP replicates (Figure 2D). Conservation of this response was also evident in the repeat experiment (Supplementary Figure 4A). During the late phase of infection, as the T_E_ population contracts, this conserved V/J signature was lost in both MT and SBP infections (Supplementary Figure 4B).

**Figure 1.**
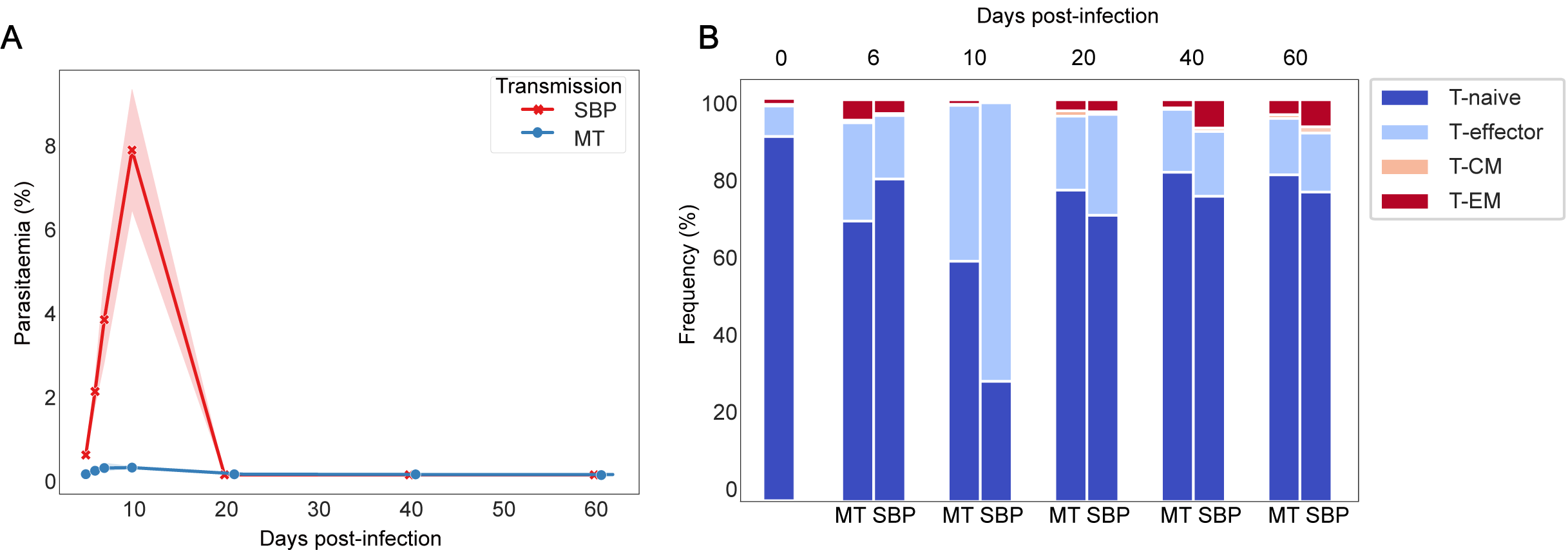
Dynamics of *P. chabaudi* infection: T-effector expansion coincides with peak parasiatemia. A) Parasitaemia of C57Bl/6 mice infected with either SBP (red) or recently MT (blue) 5 x10_5_ *P. chabaudi* parasitised erythrocytes, n=5 mice per infection type per time point, shaded area depicts 95% CI. B) Phenotypic profiling of CD4_+_ T cells as determined by FACS. Representative frequencies over the time course of infection of T-naïve CD4_+_ T cells (CD62L_+_ CD127_+_), T-effector CD4_+_ T cells (T_E_) (CD62L_-_ CD127_-_), effector memory (T_EM_) (CD44HI CD127_+_ CD62L_-_) and central memory (T_CM_) (CD44HI CD127_+_ CD62L_+_) CD4_+_T cells. Unchallenged control mice are also represented.

**Figure 2.**
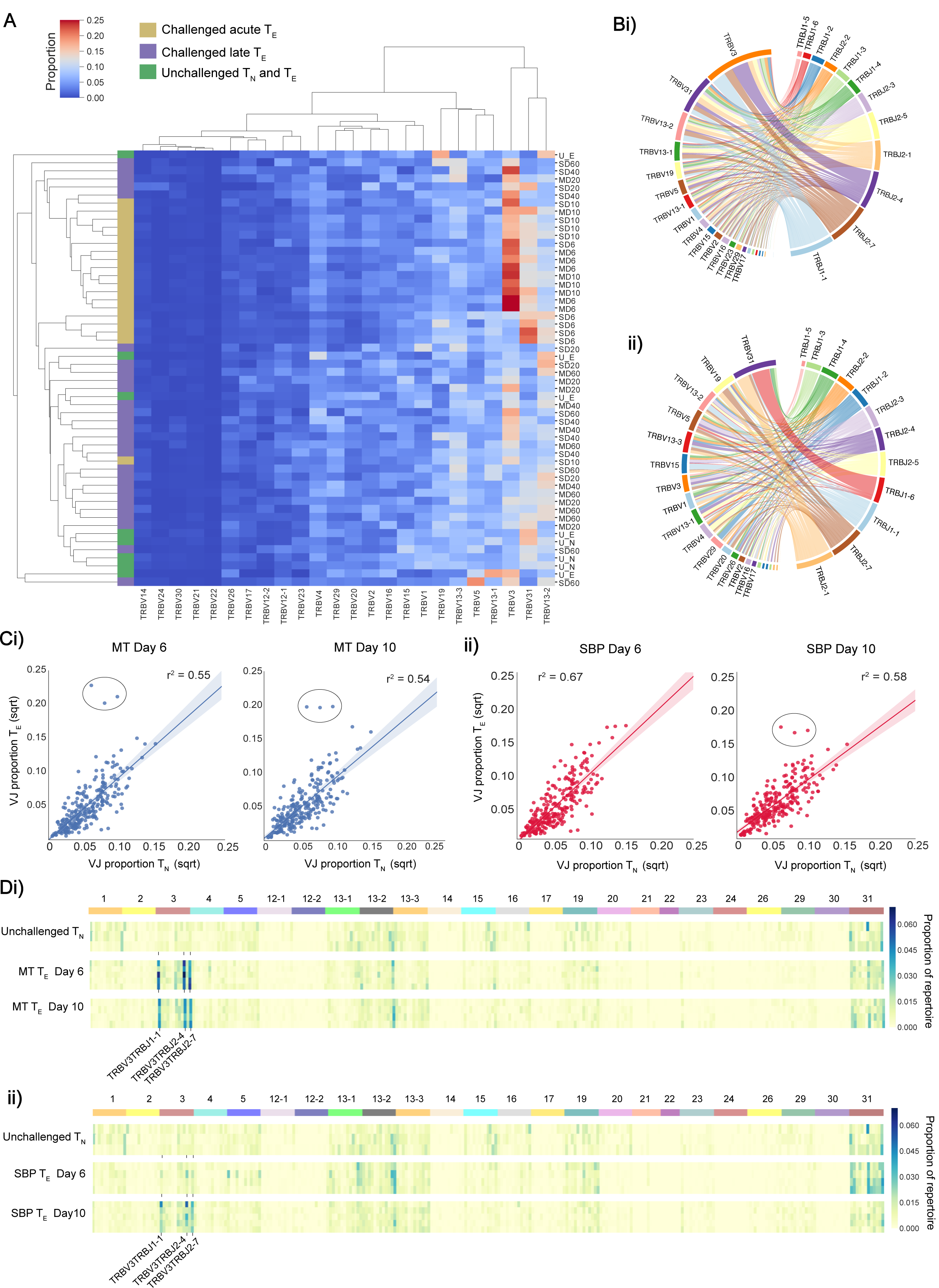
V and V/J gene usage in acute T_**E**_ repertoires: T_E_ repertoires have dominant TRBV3 gene usage during the T_E_ expansion in acute phase of a *P. chabaudi* infection. A) Clustermap displays the TRBV gene proportion of each repertoire for challenged acute T_E_ repertoires (gold), challenged late phase T_E_ repertoires (purple) and unchallenged T_N_ and T_E_ repertoires (green). B) V/J combinations from representative replicate of i) challenged T_E_ repertoire MT day 6 post-infection and ii) unchallenged T_N_ repertoire. C) Mean proportion of each V/J allelic combination in unchallenged T_N_ repertoires versus challenged T_E_ repertoires at days 6 and 10 post-infection for mice infected with (i) MT parasites (blue) and (ii) SBP parasites (red). D) Heatmaps depict proportion of each V/J allelic combination (columns) for individual replicate mice (rows) for (i) unchallenged T_N_ repertoires and acute T_E_ MT repertoires and (ii) unchallenged T_N_ repertoires and acute T_E_ SBP repertoires. Data for both days 6 and 10 post-infection are depicted. Horizontal colour bar indicates TRBV gene used in the V/J combination.

**Figure 3.**
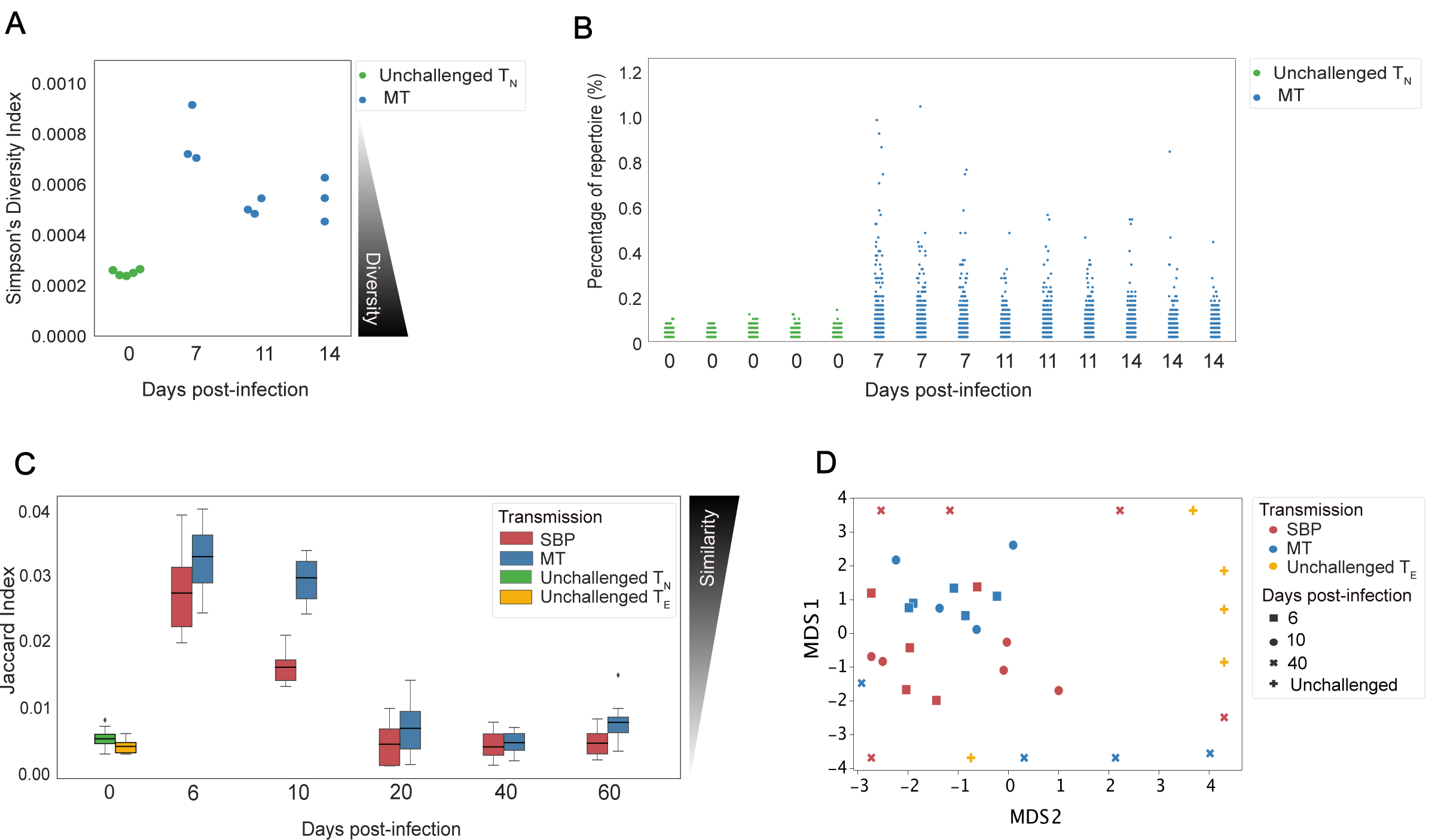
Splenic T_**E**_ repertoires elicited by *Plasmodium chabaudi* are highly diverse. A) Simpson’s Diversity Index (SI) for UMI-size matched unchallenged T_N_ (green) and acute MT T_E_ repertoires (blue). SI varies from 0 to 1, and for TCR repertoires, SI of 0 represents maximal diversity. B) Each point in the strip plot represents a clone and the percentage of repertoire they occupy, for individual replicate mice for T_N_ unchallenged repertoires (green) and MT T_E_ repertoires (blue). Data for A) and B) was normalized by down-sampling to 5000 UMI. C) T_E_ replicate repertoires are more similar to each other in the acute phase of infection for both infection types. D) Multi-dimensional-scaling (MDS) analysis using Jaccard similarity index of T_E_ repertoires for unchallenged (yellow), MT (blue) and SBP (red) repertoires. Data for C and D was normalised by down-sampling to 10_4_ reads and calculated on weighted data to include clonotype frequency.

**Figure 4.**
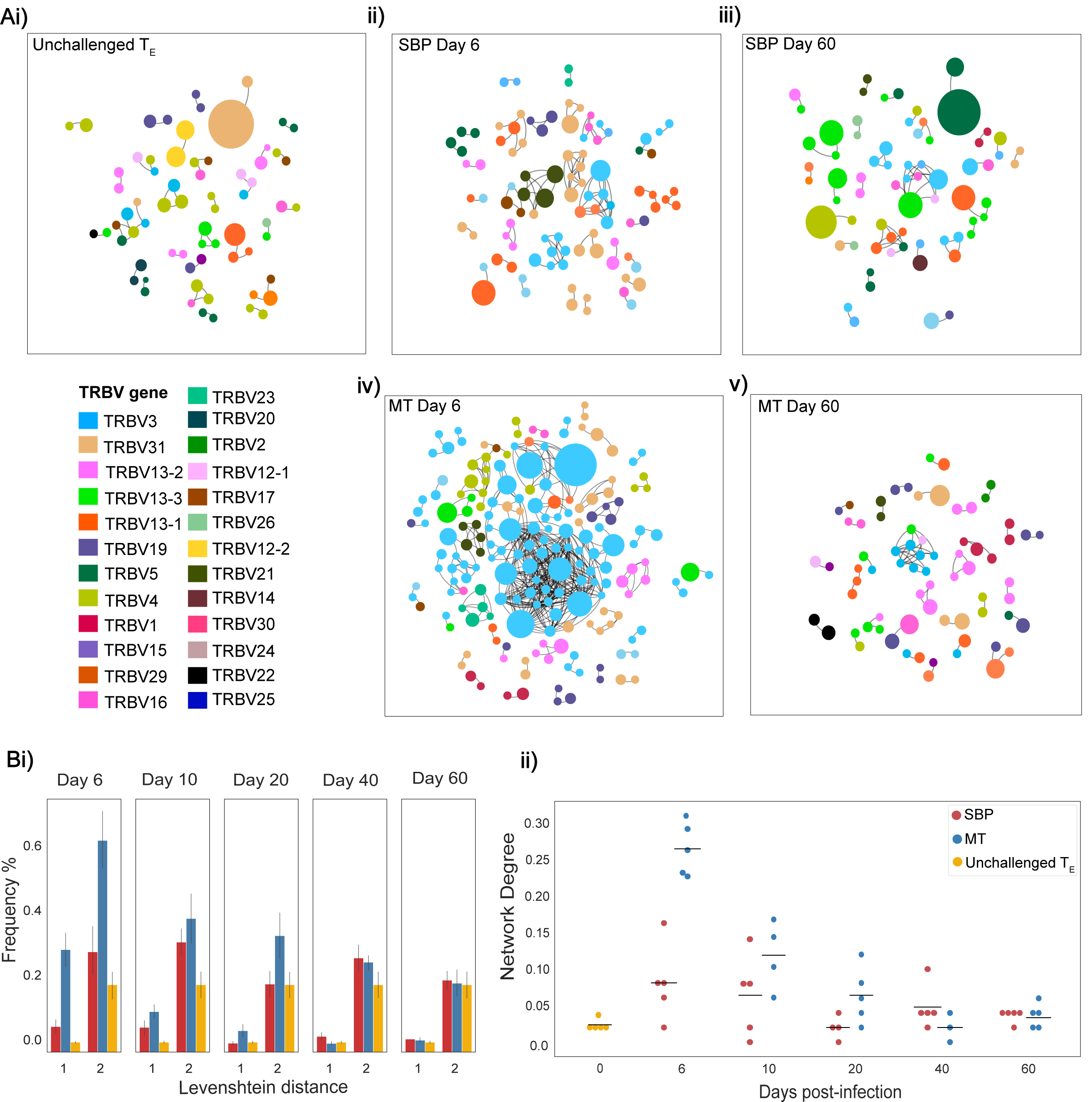
MT acute repertoires have greater clonal network connectivity. Ai-v) Networks showing the top 100 most abundant CDR3 amino acid sequences in replicate repertoires within a Levenshtein distance of 1. Each node represents a TCR clone as defined by CDR3 amino acid sequence, with node size indicating proportion of repertoire occupied by clone. Nodes are coloured according to TRBV-gene usage. An edge is drawn between nodes if within a Levenshtein distance of 1, with unconnected nodes not depicted. Bi) Frequency of individual pairwise-comparisons, between the top 100 most abundant CDR3 sequences of replicate T_E_ repertoires, that are within a Levenshtein distance of 1 and 2 and ii) network degree (mean number of edges per node) for each network of T_E_ repertoire replicates.

Examining this more precisely at the clonotype level, changes in the diversity of a TCR repertoire post pathogen exposure are indicative of the extent to which clonal expansion within a repertoire has occurred (31). To determine the diversity of repertoires during the acute CD4_+_ T_E_ expansion, repertoire diversity was calculated using Simpson’s diversity index on age-matched (32) and size-matched repertoires from the repeat experiment. This allowed us to sample an equal number of UMI-labelled cDNA molecules for the precise normalisation required for comparing diversity metrics (33). For TCRs, Simpson’s diversity index is the probability that identical TCRs (as determined by identical CDR3 amino acid sequence) will be drawn from the repertoire with two independent draws. Although unchallenged T_N_ repertoires were, as expected, significantly more diverse than challenged T_E_ repertoires (Figure 3A, p<0.01), the T_E_ repertoires after challenge were still highly polyclonal; the most abundant clone in the challenged T_E_ repertoires taking up on average only 0.72% (± 0.11%) of the repertoire compared to 0.112% (± 0.01%) in T_N_ unchallenged repertoires (Figure 3B).

### 3.2 T-Effector repertoires have greater similarity during acute infection

If there is a shared pathogen-specific response at the clonal level within the T_E_ expansion, it is expected the repertoires of replicate mice would contain conserved identical expanded antigen-specific clones, and therefore be more similar to each other than unchallenged repertoires. At the CDR3 amino acid sequence level, this may also reflect functional similarity of repertoires (33). To examine the degree of repertoire conservation between replicate mice, the Jaccard index, a similarity or ‘overlap’ metric was used, matching at the CDR3 amino acid sequence level. Over the course of infection, for both infection types, similarity between replicates is significantly altered (one-way ANOVA, MT: p<0.001, SBP: p<0.001) (Figure 3C), with replicates being more similar to each other in the acute phase of infection at days 6 and 10, than at later time-points. MT repertoires are also more similar to each other during the acute phase than SBP infections are (day 6: t=2.6, p=0.016, day 10: t=7.2, p<0.001). During the acute phase, for both infection types, replicate repertoires are more similar to each other than to unchallenged T_N_ (day 6: MT: t=15.13, p <0.001, SBP: t=11.7, p<0.001, day 10: MT: t=13.4, p<0.001, SBP: t=9.2, p<0.001) and unchallenged T_E_ repertoires (day 6: MT: t=15.1, p<0.001, SBP: t=11.7, p <0.001, day 10: MT: t=13.4, p<0.001, SBP: t=9.2, p<0.001). From day 20, effector repertoires were as dissimilar to each other as unchallenged repertoires for both infection types. Randomly sampling the same number of sequences (10_4_) to produce size-matched repertoires did not alter this pattern of results, nor did size-matching the repeat UMI data (Supplementary Figure 5). Thus, the differences in similarity are not a feature of differences in clone or read number found in replicate repertoires. Further exploration using multi-dimensional scaling (MDS) of size-matched repertoires, also indicated clustering of acute T_E_ repertoires for both infection types according to Jaccard similarity, with MT repertoires at day 6 and 10 most tightly co-clustered (Figure 3D).

**Figure 5.**
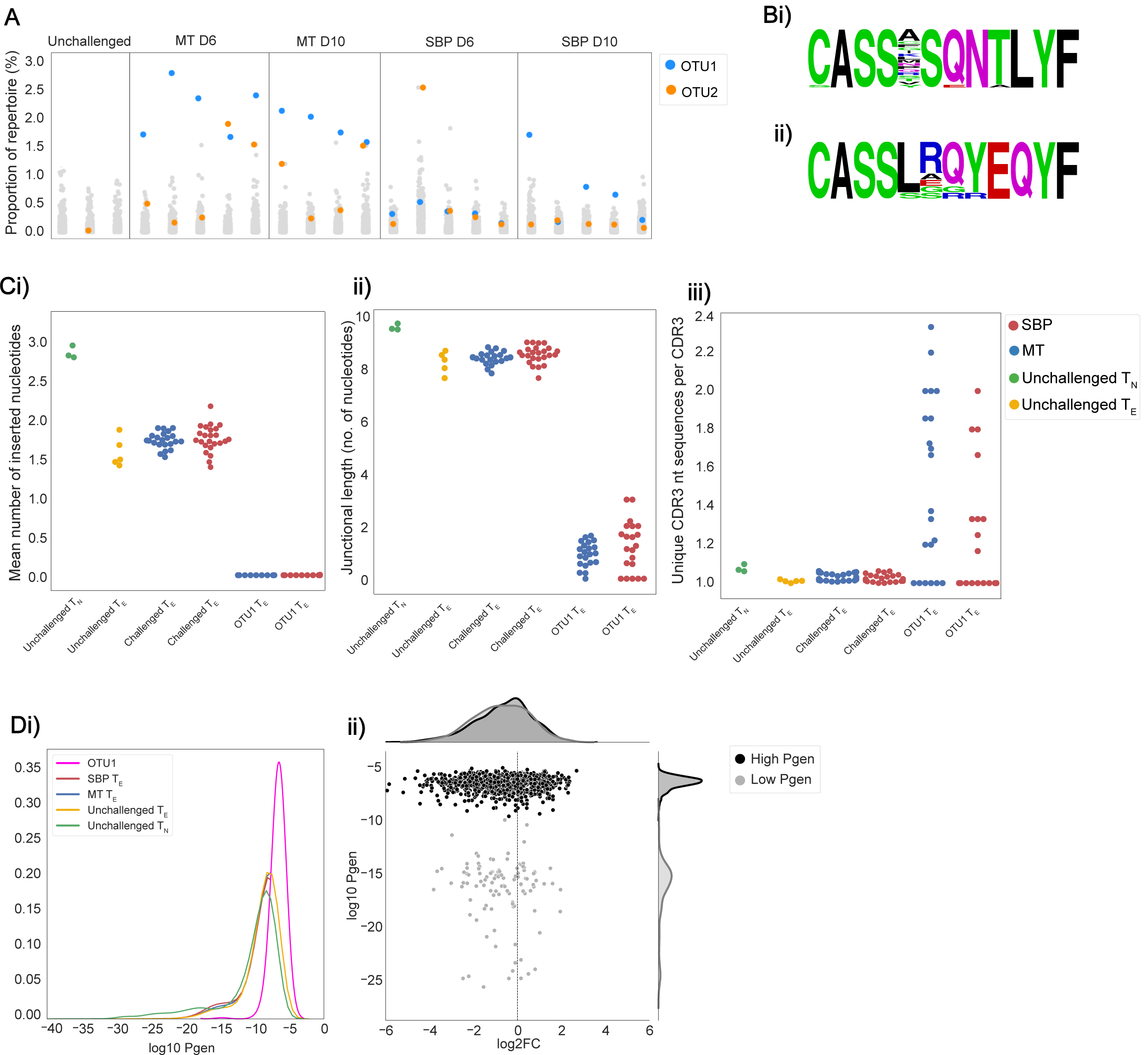
Acute MT T_**E**_ repertoires are dominated by the same cluster of clones. Repertoires were clustered using a modified Swarm algorithm, to cluster CDR3 sequences with within 1 amino-acid mismatch of each other, with identical V-gene usage. A) Strip plots display the proportion of repertoire occupied by each cluster in each individual repertoire for unchallenged T_N_ and MT and SBP T_E_ repertoires at day 6 and 10 post-infection, with OTU1 in blue and OTU2 in orange. B) Representative amino acid sequence logos of clusters OTU1 and OTU2). Ci) Mean number of inserted nucleotides of CDR3 NT sequences, ii) mean number of nucleotides lying between V and J gene segment sequences and iii) convergence – mean number of unique nucleotide sequences that encode a particular CDR3 sequence, of unchallenged T_N_ (green), unchallenged T_E_ repertoires (yellow), all MT T_E_ repertoires (blue) and all SBP T_E_ repertoires (red) and for cluster OTU1 in all MT and SBP repertoires. Di) KDE plot of the probability of generation (log10) of CDR3 nucleotide sequences for unchallenged T_N_ (green) and T_E_ repertoires (yellow), SBP (red) and MT (blue) T_E_ repertoires and CDR3 nucleotide sequences of cluster OTU1 (pink). ii) Log2 fold change of clones present in unchallenged T_N_ repertoires versus challenged T_E_ repertoires. Each point represents an individual clone, and Pgen is separated in to high (>-10) (black) and low (<-10) (grey). Log2FC was calculated on UMI normalised data.

The TCR repertoire is a dynamic network, so examining similarity solely at the clonal level can fail to take in to account the degree of extended clonal networks that may be present. Despite not undergoing somatic hypermutation, T-cell repertoires have been shown to contain networks generated by sequence similarity, with CDR3 sequence similarity and thus network connectivity increased in antigen-experienced repertoires (34,35). To explore the connectivity between CDR3s in the T_E_ repertoires, network analysis was undertaken between the top 100 most abundant clones, using Levenshtein distance. Creating networks between replicate repertoires, there was an increased frequency of CDR3 pairwise comparisons that differ by a distance of two or less for MT repertoires in the acute phase of infection (Figure 4B) compared to both SBP and unchallenged T_E_ repertoires, again highlighting the greater repertoire similarity between these replicates. Networks were also constructed individually for each repertoire, by creating an edge between unique CDR3 sequences (nodes) if they were within a Levenshtein distance of one of each other (Figure 4A). Node degree (the average number of edges per node within a network) indicated a higher degree of connectivity within individual CDR3 repertoires for both infection types in the acute stage of infection compared to unchallenged T_E_ repertoires (Mann-Whitney U, (day 6: MT: p<0.01, SBP: p=0.022, day10: MT: p<0.01, SBP: p=0.045) (Fig. 4Bii)), and also at day 20 post-infection for MT infections only (Mann-Whitney U, MT: p=0.028, SBP: p=0.38). For later time points, there was no significant difference between unchallenged and challenged network degrees detected. Between infection types, at days 6, 10 and 20 post-infection, MT repertoires also tended to have a higher node degree than SBP repertoires at the equivalent time-points, although day 10 was non-significant (Mann-Whitney U, day 6: p<0.01, day 10: p=0.088, day 20: p=0.039).

### 3.3 The same ‘public’ cluster is most dominant in the majority of acute T-effector repertoires

Although none of the most dominant clone of each repertoire was identical between repertoires, given the increased connectivity in challenged repertoires, and the knowledge that TCRs recognizing the same antigen typically have a high global similarity to each other (35,36), we clustered the CDR3 sequences of individual repertoires within one amino acid mismatch of each other using a modified Swarm algorithm (24). This identified two clusters of highly similar CDR3 sequences, hereafter referred to as OTU1 and OTU2, that were conserved across replicates in the acute phase of infection. For MT samples in the acute phase of infection, OTU1 was the most abundant cluster in the majority of replicates at both day 6 and day 10 post-infection (Figure 5A, 5Bi). For SBP infections, OTU1 was not prominent at day 6, but became the most abundant cluster in 3 out of the 5 replicates by day 10 (Figure 5A). CDR3s in OTU1 were encoded by TRBV3-TRBJ2-4. OTU2 was also identified as a conserved cluster across replicates, but was only dominant in some (Figure 5A). The Swarm algorithm was applied separately on the repeat experiment, and a near-identical cluster to OTU1 was again found to be expanded and dominant in a majority of challenged repertoires at day 7 and 11 post-infection (Supplementary Figure 6B). Both OTU1 and OTU2 were also observed in the analyses of publicly available RNA-seq data sets for *P. chabaudi (AS)* and *P. chabaudi (CB)* (accession E-ERAD-221 and E-ERAD-289 respectively, Supplementary Figure 6C), showing a similar temporal pattern of expansion in the acute phase of infection. We also applied the recently published Gliph2 algorithm (25) to our data. Gliph2 is designed to identify TCRs recognizing the same antigen by clustering sequences with enriched amino acid motifs in the high-contact-probability region of CDR3β (IMGT positions 107-116). It identified significant clusters that corresponded to both OTU1 and OTU2 in the acute phase of MT infections, as well several other clusters which were not as dominant nor as well-conserved in their response (Supplementary Table 1, Supplementary Figure 6A). Given the conserved nature of OTU1 between individual mice, we hypothesised that the CDR3 sequences it contains would share similar properties with other known ‘public’ CDR3 sequences, defined simply as TCR clones shared by different individuals. Public TCR clones have been detected in numerous T-cell responses in multiple species, and although their functional significance remains unknown, they have been shown to be expanded in response to antigenic stimulation (37), viral infection (38,39) and associated with regulatory self-immunity (40). In some previous studies, public sequences have been shown to have minimal alterations to germline V, D and J gene sequences. In agreement with this, we found fewer recombination events in the CDR3 sequences in OTU1, with the mean number of randomly inserted nucleotides in the CDR3 sequences in these clusters significantly lower than that for CDR3 sequences in both challenged (t=−61.5, p<0.001) and unchallenged repertoires (t=−58.7, p<0.001) (Figure 5Ci). The mean number of nucleotides lying between the V and J gene segment sequences was also significantly lower (unchallenged: t=−22.1, p=0.002, challenged: t=−83.7, p<0.001) (Figure 5Cii). This cluster also showed a greater degree of convergent recombination – considered an important mechanism of public TCR generation (37,41) – with a higher average number of unique CDR3 nucleotide sequences that code for the same CDR3 amino acid sequence (Figure 5Ciii) compared to CDR3 sequences in unchallenged (t=4.4, p<0.001) and challenged (t=5.3, p<0.001) repertoires. Consequently, CDR3 amino acid sequences in OTU1 had a higher probability of generation (Pgen) (27) compared to all CDR3 sequences in unchallenged T_N_ repertoires and challenged repertoires (Figure 5Di). In a scenario of non-specific polyclonal expansion, a higher Pgen could indicate sharing and detection of this cluster incidentally due to higher abundance in the pre-selection pool, rather than as a result of common specificity (convergent selection) or function (42). However, the cluster was either not found or was present at a very low level in unchallenged T_N_ and T_E_ repertoires, and our use of UMI-corrected data for the repeat experiment together with gating strategy, confirmed that the CDR3s within this cluster are clonally expanded. Further, a large proportion of CDR3s with a similar high Pgen in unchallenged T_N_ populations, were either not present in T_E_ repertoires, or present but not expanded, demonstrating that the public cluster is likely to be pathogen-associated (Figure 5Dii).

**Figure 6.**
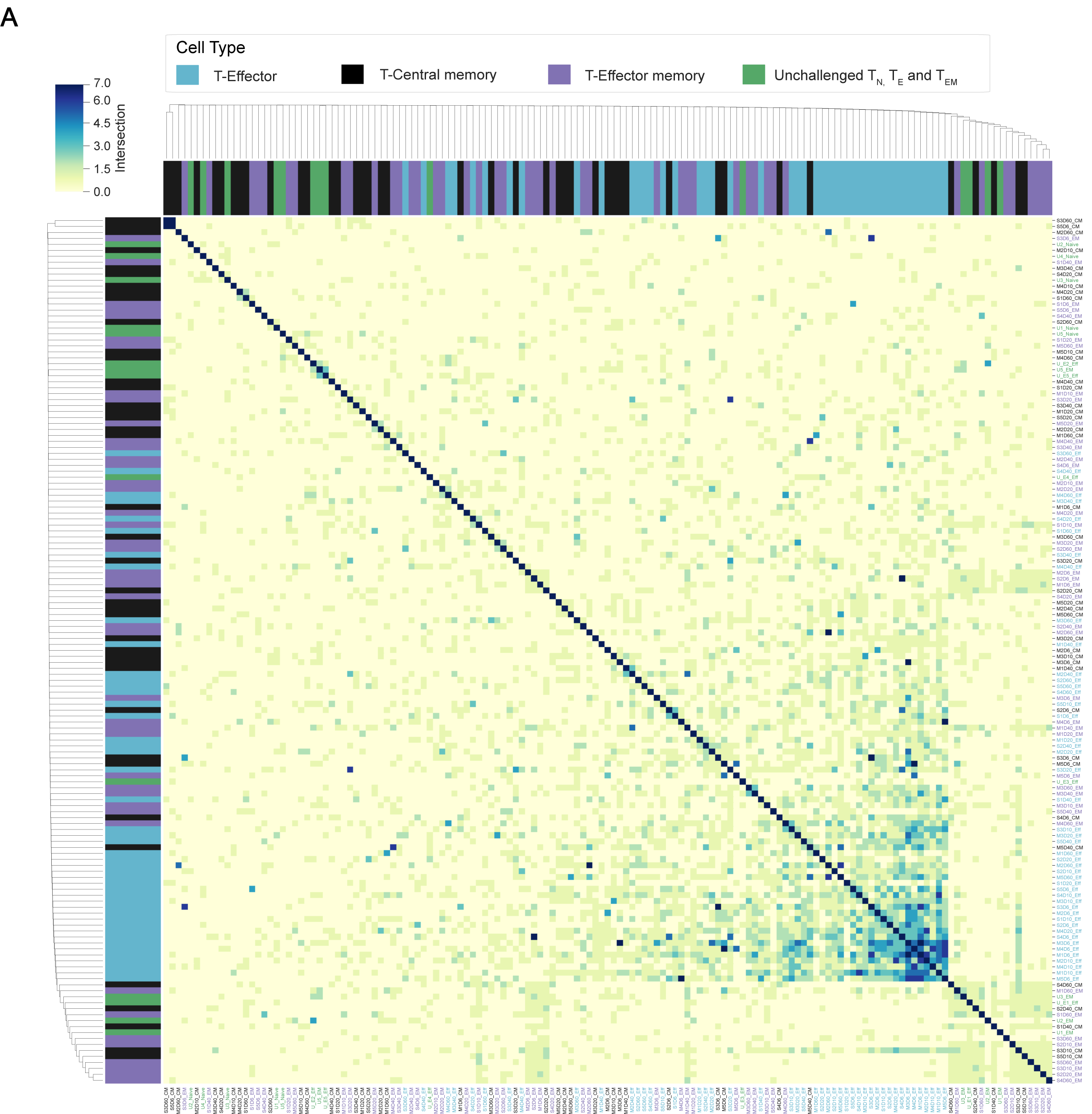
Splenic memory responses are private responses: A) Cluster map depicts the pairwise number of overlapping CDR3 amino acid sequences in the top 100 CDR3 sequences of individual unweighted challenged T_E_ (blue), T_CM_ (black) and T_EM_ (purple) repertoires, and unchallenged T_N_, T_E_ and T_EM_ repertoires (green).

To determine if the CDR3 sequences in OTU1 have been shown to expand in response to other antigenic stimuli in C57Bl/6 mice, we analysed publicly available splenic CD4_+_ TCRβ repertoire data (40) from unchallenged mice and mice immunised with either OVA or CFA and OVA. Although detectable, the proportion of OTU1 did not differ between unchallenged and immunised mice, indicating the expansion seen is not simply an innate response to inflammation. TCR repertoire data was also extracted from publicly available whole blood RNA-seq data from C57Bl/6 mice challenged with a variety of pathogens; *Toxoplasma gondii*, influenza A virus, murine cytomegalovirus, respiratory syncytial virus, *Candida albicans, Listeria monocytogenes, Burkholderia pseudomallei*, and House dust mite allergen (43). Of the 9 pathogens examined at the peak of the murine response, only *P. chabaudi* elicited an expansion in TRBV3 gene usage (Supplementary Figure 7). As such, OTU1 was not found to be expanded by any of the other pathogens examined, supporting the probable specificity of this response.

**Figure 7.**
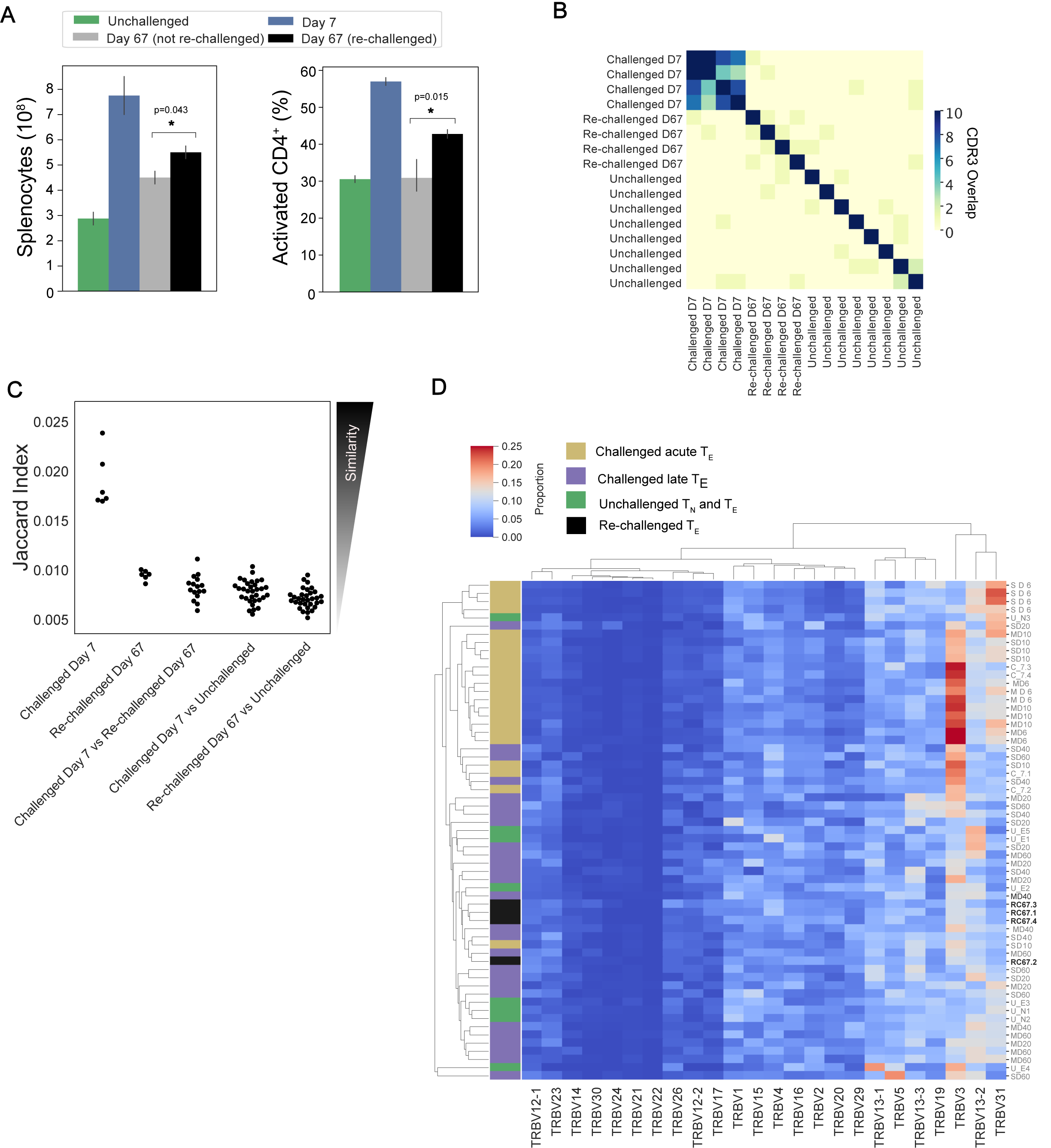
No conservation of response following homologous re-challenge: A) Splenocyte numbers and frequency of activated CD4_+_T cells (CD44HI) of unchallenged mice (green), day 7 post-primary infection (blue), day 67 post primary infection (grey, not re-challenged) and 7 days following re-challenge (black, re-challenged day 67). B) Pairwise overlap of the top 100 most abundant CDR3 amino acid sequences of repertoires for the re-challenge experiment. C) Jaccard (similarity) index of replicate repertoires (normalized by down-sampling to 5000 UMI) and D) Clustermap displays the TRBV gene proportion of each repertoire for challenged acute T_E_ repertoires (gold), challenged late phase T_E_ repertoires (purple) and unchallenged T_N_ and T_E_ repertoires (green) from the first experiment, with re-challenged T_E_ repertoires (black) now also included.

### 3.4 Splenic memory and re-challenged repertoires are private responses

Splenic CD4_+_ T_EM_ and T_CM_ populations have been shown to expand following a *P. chabaudi* infection (8). To determine if a conserved splenic memory response was also generated, the splenic T_EM_ and T_CM_ populations from the first experiment were sequenced, for both infection types. All similarity and CDR3 overlap analyses were conducted unweighted, to avoid biases from effector clonal expansion. Little to no sharing was found between either the T_EM_ or T_CM_ replicate repertoires themselves for either infection type, nor between the T_EM_, T_CM_ and T_E_ populations (Figure 6A). Clustering the memory repertoires within one amino acid mismatch using the same modified Swarm algorithm (24) did not improve the similarity between memory replicates, indicating these responses are ‘private’ to each individual at this sequencing depth.

As a conserved response was not detected within memory populations, a homologous rechallenge infection was undertaken to determine if priming of identified clusters of interest had occurred. Mice were re-challenged at day 60 post-primary infection, and then sacrificed at day 67 (day 7 post re-challenge). Following re-challenge, mice did not develop a detectable parasitaemia, but a splenic expansion of an activated effector population was evident, though not as marked as following primary challenge (Figure 7A). Although TRBV3 was again confirmed to be expanded 7 days after primary challenge, no re-expansion of TRBV3 was observed following re-challenge, with T_E_ repertoires clustering with later time-points rather than acute repertoires (Figure 7D). At the clonal level, rechallenged repertoires were found to be as dissimilar to each other as unchallenged repertoires (Figure 7C), with little to no sharing of the 100 most abundant CDR3 amino acid sequences (Figure 7B).

### 3.5 Activated TRBV3_**+**_ cells have diverse transcriptional phenotypes

To further our understanding of the phenotype of the conserved TRBV3 response observed during the acute phase of a *P. chabaudi* infection, we undertook single-cell RNA sequencing of FACS sorted CD3_+_ CD4_+_ splenocytes from two mice at day 7 post-infection with MT parasites. We sequenced these cells using the 10x Genomics Chromium platform, using the V(D)J enrichment protocol to obtain paired α/β TCR data for each cell. After quality control steps, we obtained expression profiles for 1491 and 1485 CD4_+_ T cells from each mouse respectively (2976 total). Following dimensionality reduction by principal component analysis (PCA), we undertook graph-based clustering (29) and visualised resulting populations using uniform manifold approximation and projection (UMAP) (Figure 8A). We identified seven discrete transcriptional clusters, with cells from both mice evenly distributed across all dominant clusters (Supplementary Figure 8). Four of these clusters, denoted as clusters 1, 2, 3 and 4, were classified overall as naïve on the basis of canonical markers (*Sell, Il7r*) (Figure 8B, Supplementary Figure 9). Differential expression between all seven clusters indicated that clusters 2 and 3 were distinguishable by markers indicative of early T-cell activation, including *cd69, ltm2a, Zfp36* and *ler2* for cluster 3, and type I interferon (IFN) response genes (*Gbp2, Ifit1, Ifit3, Isg15*, and *Socs1*) (44) for cluster 2 (Supplementary Figure 10), whilst cluster 1 expressed a more definitive naïve phenotype (*lef1, Ccr7*). We also identified three clusters discrete from the naïve group: cluster 6 showed a typical Treg transcriptional phenotype with expression of *Foxp3* and *Il2ra*, whilst clusters 5 and 7 were both identified as activated effector populations (*Sell-, Il7r-*). Cluster 7 expressed genes associated with a pro-inflammatory T_H_1 signature, including *Tbx21, Ifng, Gzmb, Cxcr6, Ccl5, NKg7* and *Bhlhe40* (45). In contrast, cluster 5 was *Ifng-* and appeared to be a more diverse helper population, which predominantly included cells expressing genes associated with a T_FH_ phenotype (*Cxcr5, Icos, Bcl6, Il21, Pdcd1)* as well as those with a T_H_2 phenotype (*Gata3, Ccr4*) (Figure 8C). Utilising the TCR data, we confirm that the effector response is highly polyclonal: of the 581 effector cells present in clusters 5 and 7, only 3.6% (21/581) are clonally expanded (>1 cell with identical paired TCRα and TCRβ aa sequence) (Figure 8D), and no clone had more than three copies present. We again confirm that TRBV3 is the most dominant TRBV gene encoding the TCRβ chain of activated effector populations, and show that out of all the clusters, cluster 7 has the highest proportion of TRBV3 (Figure 8E). Cells that are TRBV3_+_ do not however form a distinct cluster and instead display diverse phenotypes across all clusters. No TRAV gene is over-represented in a particular cluster (Supplementary Figure 8B). Of the clonotypes previously identified in the TCRβ bulk data in OTU1 and OTU2, two are present in the single-cell data set, both within the predominantly T_FH_ cluster 5. TCRβ nucleotide sequences displayed diverse probabilities of generation across all clusters (Supplementary Figure 11).

**Figure 8.**
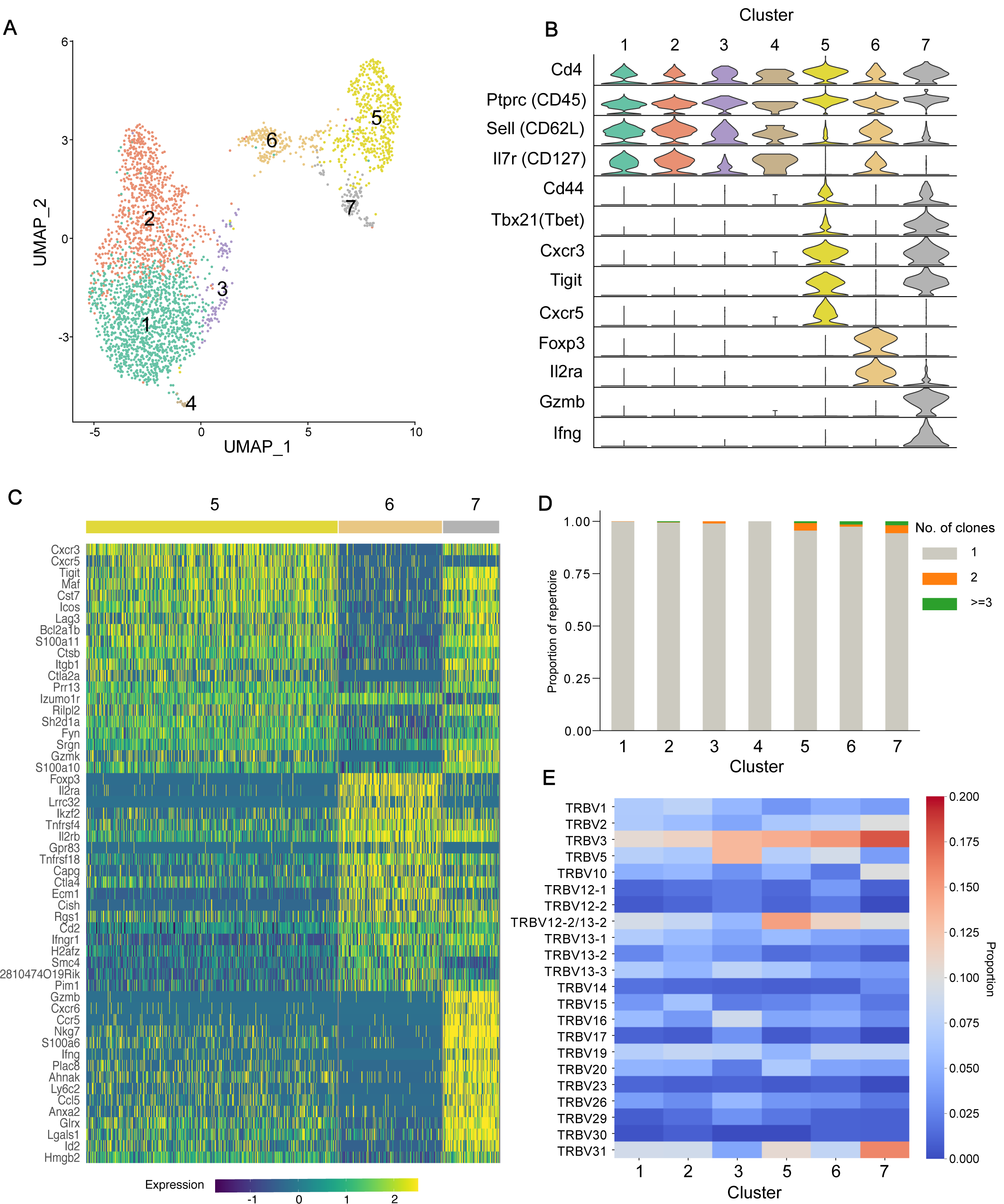
Activated TRBV3_**+**_ cells have diverse transcriptional phenotypes: A) Two-dimensional UMAP visualization of CD4_+_ splenocytes from 2 challenged mice, at day 7 post-infection, as per Seurat cluster assignment. B) Violin plots display normalized expression of major canonical T-cell markers per cluster. C) Top 20 differentially expressed genes that define clusters 5, 6 and 7. D) Proportion of each cluster that is clonally expanded (>1 cell with identical paired TCRα and TCRβ aa sequences). E) Heatmap displays proportion of TRBV gene per cluster TCR repertoire.

**Figure 9:**
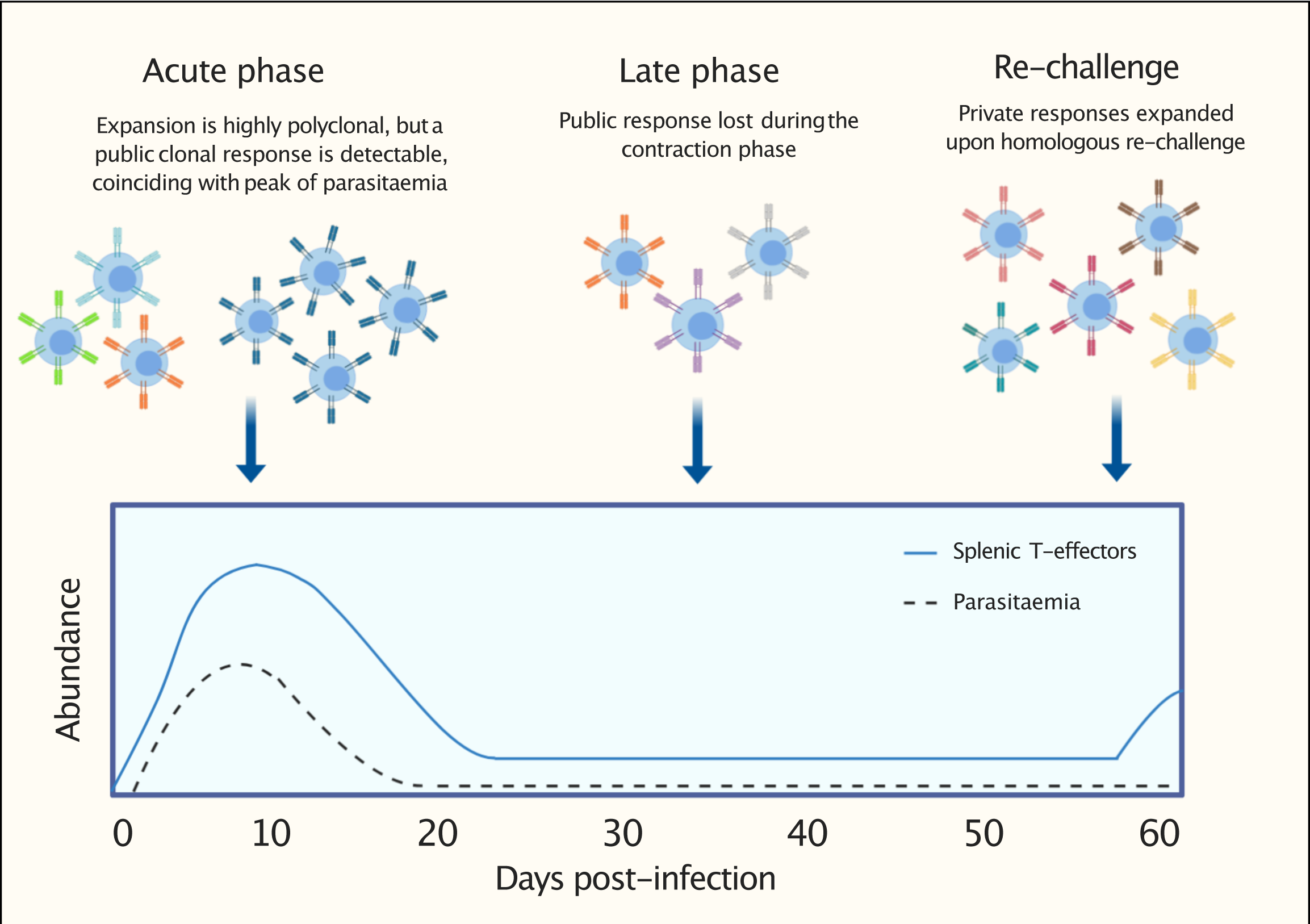
Graphical representation to summarise findings: following a first *P. chabaudi* infection, within a highly polyclonal T-effector expansion, a public TCRβ clonal response encoded by TRBV3 is detectable. This conserved response contracts during the late phase of infection and is not re-expanded upon homologous re-challenge. This response is consistently found to be a hallmark of a first *P. chabaudi* infection.

## 4. Discussion

CD4_+_ T-cells play a critical role in the immune response against the pathological blood-stage of malaria (reviewed in (6,7)). However there is a lack of deep mechanistic understanding regarding the development of T-cell mediated immunity against *Plasmodium* (7). To our knowledge, this is the first study to use bulk TCR deep-sequencing to examine the composition of CD4_+_ T-cell repertoires induced by a *Plasmodium* infection. We found that in both the MT and SBP infection models, which show differential control of parasite growth and degrees of immunopathology (8,16), T_E_ repertoires elicited upon infection are highly diverse and polyclonal. Despite the 10-fold increase in splenic T_E_ cell numbers, the V/J gene usage frequencies of acute challenged T_E_ repertoires are strongly positively correlated with that of unchallenged T_N_ repertoires, indicating a highly polyclonal broad expansion of the naïve repertoire, consistent with the massive cellular expansion seen. This polyclonality is confirmed in our single-cell RNA-seq data. The degree of correlation between VJ usage frequencies likely indicates a degree of non-specific expansion, although a highly heterogenous response to the vast number of potential antigens expressed by the parasite, argued as a potential cause for the preponderance seen in *P. falciparum* IgG responses (46), cannot be ruled out. However, within this highly polyclonal effector proliferation, a strong and oligo-clonal conserved response is observed in the bulk TCR data following a first infection. Here, repertoires are skewed to TRBV3 gene usage, have a higher degree of clonal sharing, and show increased amino acid sequence similarity of the CDR3 region between dominant clones. Thus, we demonstrate that despite the antigenic complexity of *P. chabaudi*, T_E_ repertoires bear the hallmarks of a specific response (35,36), mirroring that observed with less antigenically diverse organisms (47–49). This conserved response is evident in both infection models, although it is delayed and less marked in mice infected with SBP parasites. The similarities suggest that the drivers of this response are likely to be common to both MT and SBP parasites. Sequencing of total parasite RNA for both MT and SBP parasites undertaken by Spence et al (2013), indicated the parasites’ *cir* gene family – believed to encode a large set of variable antigens displayed on parasitized erythrocytes – was differentially expressed between the two infection models. MT parasites upregulated expression of many genes within this family equally, re-setting broad expansion of the antigenic repertoire, whilst SBP selected for a limited but dominant *cir* expression. Our results are therefore inconsistent with proteins encoded by *cir* genes driving the detected conserved response, which would instead have been expected to result in a more dominant response in SBP infections. We hypothesise that the differences observed in the timing and magnitude of the TRBV3-restricted shared response seen between the two infection models are a result of the systemic inflammation induced by SBP parasites (8) disrupting or delaying the formation of an appropriate T-cell response. Future work to determine MHC-presentation pathway and ligands or peptides involved is ultimately required to determine what specific parasite epitope elicits the conserved response.

The single cell RNA-seq data confirms the dominance of TRBV3 in activated effector populations and reveals that TRBV3_+_ cells in the acute phase of a first infection have diverse phenotypes. We therefore show that the conserved response seen does not represent a discrete innate cell population but is instead part of an adaptive immune process against the parasite. The transcriptional phenotypes of activated effector cells are in agreement with those previously observed in the acute phase of a *P. chabaudi* infection (45) and indicate a large cluster of predominantly follicular helper cells as well as a distinct *Ifng*_+_ T_H_1 population. Both types of response have been shown to arise simultaneously during the acute phase (45) and to be essential in controlling blood-stage *P. chabaudi*; T_H_1 responses are required for initial control of acute parasitaemia (50,51) and T_FH_ are crucial for generating antibody-mediated immunity and controlling chronic infection (52,53). The presence of OTU1 and OTU2 in a predominantly T_FH_ cluster, suggests that they may play a role in guiding the developing humoral response against the parasite. A single naïve CD4_+_ T-cell has been demonstrated to be able to give rise to clones with different cell fates (45) so, as only one of each clonotype was captured with the single cell sequencing depth used, we cannot firmly conclude that the transcriptional phenotypes identified here would broadly reflect the conserved TCRs identified in the bulk data. Nevertheless, the TCR of a cell has been shown to impart a strong preference for either a T_H_1 or a T_FH_ fate, with longer dwell time between peptide-MHC:TCR biasing towards T_FH_ and GC-T_FH_ responses (54). Therefore, if driven by the same epitope, we would expect the conserved response to have a similar phenotype (55).

Public TCR sequences are shared between multiple individuals either due to biases in V(D)J recombination, and/or convergent selection by a common antigen (42,47). The CDR3 amino acid sequences in the most dominant and conserved cluster detected (OTU1) had similar features to other previously observed public TCR responses, including a reduced number of recombination events, a greater degree of recombinant convergence and subsequent higher probability of generation (PGen). The degree of clonal expansion observed in OTU1 sequences was greater than many other sequences of equal or higher PGen, indicating that these clones were truly expanded and not simply found to be of high frequency as a result of recombinational biases. We also show that PGen does not determine T-cell fate, in agreement with Sethna *et al* (2018) who demonstrated that the ability of a TCR to respond to a particular epitope, was not strongly correlated with its generation probability. It has been hypothesised that public TCR responses may provide rapid cross-reactive immunity (56) to cope with diverse antigenic challenge, allowing time for more specific private responses to develop (37,57). Thus, during a *P. chabaudi infection*, a public response that is mobilised rapidly due to high Pgen and/or a higher chance of positive selection if cross-reactive, may act as a first line of defence against the parasite before more specific responses become effective. In agreement with this, OTU1 appears to be temporally associated with enhanced control of parasitaemia. It arises earlier and is more dominant in MT infections compared with SBP, where rapid parasite growth is observed alongside a delayed and less marked conserved response. Mice infected with MT parasites also show reduced disease severity (8). Despite these positive associations, whether this conserved response is truly beneficial to the host, remains unknown. There are reports of public TCR responses being implicated in self-related immunity (40), and in *P. berghei*, the presence of conserved pathogenic CD8_+_ T-cells has been used to predict cerebral malaria (11).

No conserved response was evident in memory populations or upon homologous rechallenge. One explanation for this is that conserved TCRs identified in the effector population lack the avidity required to enter the memory pool (58). Alternatively, Opata et al (*2015*) suggested that the highly active CD62L_-_ CD127_-_ T_E_ population (as captured by our gating strategy) in a *P. chabaudi* infection may be terminally differentiated, and it is an earlier T_E_ population (CD127_-_ CD62L_hi_) that contains memory precursors. Thus, with the gating strategy used to capture maximum effector function, this population may have been excluded. However, a separate conserved memory response would have been detectable through similarity indices and clustering of memory repertoires. This was not evident, indicating splenic memory responses are private to each individual. This is consistent with only private responses being expanded upon homologous re-challenge, where mice were protected from re-infection. Such private responses may target the same or multiple *Plasmodium* antigenic targets.

To have found a near-identical expanded T_E_ cluster in independent repeat experiments, and in publicly available *P. chabaudi* (AS and CB) RNA-seq data sets, demonstrates the TCR signature is consistently expanded in multiple individuals in response to a first *Plasmodium chabaudi* infection. TRBV3 was not expanded at peak infection for any of the other eight pathogens examined and searches of annotated TCR sequence databases have not revealed any other known specificities for any of the CDR3 amino acid sequences in OTU1 or OTU2 (48,59). TCR repertoire and RNA-seq data sets following infection with other *Plasmodium* species are currently unavailable, but as these are generated, the specificity of this response to the parasite and whether a similar response is elicited in a first infection by other *Plasmodium* species including *P. falciparum* will become evident.

Future studies are required to determine whether the conserved response plays a critical role in protection during a first infection. MT infections develop chronic recrudescing infection, so the response does not fully clear infection, although it is temporally associated with control of parasite growth and reduced disease severity (8). Whether entirely favourable to the host or not, we hypothesise the response allows time for mechanisms that govern the formation of more specific private responses and subsequent immunity to the parasite to develop. Mice are known to develop highly effective strain-specific anti-parasite immunity after a single malaria episode, whilst this takes years to develop in people. Therefore, if some degree of first line protection were shown, such a response could be a novel target to promote in malaria naïve individuals through vaccination, providing initial cover whilst more specific but slower private responses develop, conveying higher levels of protection. Even if only partially protective, the response is conserved between individuals and receptor sequences have a high probability of generation. Targeting these is predicted therefore to increase vaccine success rate (27,60).

In summary, we have demonstrated that a conserved TCRβ signature encoded by TRBV3 is consistently expanded in response to a first *Plasmodium chabaudi* infection (Figure 9). In contrast, memory formation and re-challenged repertoires appear to be private responses to the individual. Understanding the antigenic drivers and contribution to protection (or pathogenesis) of this conserved signature, that is consistently a hallmark of a first infection, is ultimately required to determine if it should be promoted or mitigated for malaria therapeutic purposes.

## 5. Conflict of Interest

The authors declare that the research was conducted in the absence of any commercial or financial relationships that could be construed as a potential conflict of interest.

## 6. Author Contributions

NS, GC and CS designed the project, analysed the data, and wrote the manuscript. Experimental work was carried out by NS, GC, WN, JM, PS and JT. All authors reviewed the manuscript.

## 7. Funding

This work was made possible by a studentship from the Wellcome Trust to NS (204511/Z/16/A) and Wellcome Trust ISSF funding to GC.

## 8. Acknowledgments

This work has made use of the resources provided by the Edinburgh Computer and Data Facility (ECDF) (http://www.ecdf.ed.ac.uk/). The authors thank Dr Martin Waterfall of the IIIR Flow Cytometry Facility for help with cell sorting. Bulk TCR sequencing was carried out by Edinburgh Genomics at the University of Edinburgh. Edinburgh Genomics is partly supported through core grants from NERC (R8/H10/56), MRC (MR/K001744/1) and BBSRC (BB/J004243/1). Single-cell sequencing was carried out by the IGMM sequencing facility at the University of Edinburgh.

## Notes

### Competing Interest Statement

The authors have declared no competing interest.

## References

1. Ryg-Cornejo V, Ly A, Hansen DS. Immunological processes underlying the slow acquisition of humoral immunity to malaria. Parasitology (2016) 143:1–9. doi: 10.1017/S0031182015001705

2. Gupta S, Snow RW, Donnelly CA, Marsh K, Newbold C. Immunity to non-cerebral severe malaria is acquired after one or two infections. Nat Med (1999) 5:340–343. doi: 10.1038/6560

3. Gonçalves BP, Huang C-Y, Morrison R, Holte S, Kabyemela E, Prevots DR, Fried M, Duffy PE. Parasite Burden and Severity of Malaria in Tanzanian Children. N Engl J Med (2014) 370:1799–1808. doi: 10.1056/NEJMoa1303944

4. Ly A, Hansen DS. Development of B cell memory in malaria. Front Immunol (2019) 10:1–11. doi: 10.3389/fimmu.2019.00559

5. Doolan DL, Dobaño C, Baird JK. Acquired immunity to Malaria. Clin Microbiol Rev (2009) 22:13–36. doi: 10.1128/CMR.00025-08

6. Nlinwe ON, Kusi KA, Adu B, Sedegah M. T-cell responses against Malaria: Effect of parasite antigen diversity and relevance for vaccine development. Vaccine (2018) 36:2237–2242. doi: 10.1016/j.vaccine.2018.03.023

7. Kurup SP, Butler NS, Harty JT. T cell-mediated immunity to malaria. Nat Rev Immunol (2019) doi: 10.1038/s41577-019-0158-z

8. Spence PJ, Jarra W, Lévy P, Reid AJ, Chappell L, Brugat T, Sanders M, Berriman M, Langhorne J. Vector transmission regulates immune control of Plasmodium virulence. Nature (2013) 498:228–31. doi: 10.1038/nature12231

9. Opata MM, Stephens R. Chronic Plasmodium chabaudi infection generates CD4 memory T cells with increased T cell receptor sensitivity but poor secondary expansion and increased apoptosis. Infect Immun (2017) 85:1–16. doi: 10.1128/IAI.00774-16

10. del Portillo HA, Ferrer M, Brugat T, Martin-Jaular L, Langhorne J, Lacerda MVG. The role of the spleen in malaria. Cell Microbiol (2012) 14:343–355. doi: 10.1111/j.1462-5822.2011.01741.x

11. Mariotti-Ferrandiz E, Pham HP, Dulauroy S, Gorgette O, Klatzmann D, Cazenave PA, Pied S, Six A. A TCRB Repertoire Signature Can Predict Experimental Cerebral Malaria. PLoS One (2016) 11:1–17. doi: 10.1371/journal.pone.0147871

12. Ndungu FM, Sanni L, Urban B, Stephens R, Newbold CI, Marsh K, Langhorne J. CD4 T Cells from Malaria-Nonexposed Individuals Respond to the CD36-Binding Domain of Plasmodium falciparum Erythrocyte Membrane Protein-1 via an MHC Class II-TCR-Independent Pathway. J Immunol (2006) 176:5504–5512. doi: 10.4049/jimmunol.176.9.5504

13. Scholzen A, Mittag D, Rogerson SJ, Cooke BM, Plebanski M. Plasmodium falciparum – Mediated Induction of Human CD25 hi Foxp3 hi CD4 T Cells Is Independent of Direct TCR Stimulation and Requires IL-2, IL-10 and TGF b. PLoS Pathog (2009) 5:26–32. doi: 10.1371/journal.ppat.1000543

14. Wipasa J, Okell L, Sakkhachornphop S, Suphavilai C, Chawansuntati K, Liewsaree W, Hafalla JCR, Riley EM. Short-lived IFN-γ effector responses, but long-lived IL-10 memory responses, to malaria in an area of low malaria endemicity. PLoS Pathog (2011) 7: doi: 10.1371/journal.ppat.1001281

15. Bradley P, Thomas PG. Using T Cell Receptor Repertoires to Understand the Principles of Adaptive Immune Recognition. Annu Rev Immunol (2019) 37: doi: 10.1146/annurev-immunol-042718-041757

16. Spence PJ, Brugat T, Langhorne J. Mosquitoes Reset Malaria Parasites. PLoS Pathog (2015) 11:10–15. doi: 10.1371/journal.ppat.1004987

17. Marr EJ, Milne RM, Anar B, Girling G, Schwach F, Mooney JP, Nahrendorf W, Spence PJ, Cunningham D, Baker DA, et al. An enhanced toolkit for the generation of knockout and marker-free fluorescent Plasmodium chabaudi. Wellcome Open Res (2020) 5:1–20. doi: 10.12688/wellcomeopenres.15587.1

18. Lewis M, Pfeil J, Mueller AK. Continuous oral chloroquine as a novel route for Plasmodium prophylaxis and cure in experimental murine models. BMC Res Notes (2011) 4:262. doi: 10.1186/1756-0500-4-262

19. Kivioja T, Vähärautio A, Karlsson K, Bonke M, Enge M, Linnarsson S, Taipale J. Counting absolute numbers of molecules using unique molecular identifiers. Nat Methods (2012) 9:72–74. doi: 10.1038/nmeth.1778

20. Shugay M, Britanova O V, Merzlyak EM, Turchaninova MA, Mamedov IZ, Tuganbaev TR, Bolotin DA, Staroverov DB, Putintseva E V, Plevova K, et al. Towards error-free profiling of immune repertoires. (2014) 11:6–10. doi: 10.1038/nmeth.2960

21. Bolotin DA, Poslavsky S, Mitrophanov I, Shugay M, Mamedov IZ, Putintseva E V, Chudakov DM. MiXCR : software for comprehensive adaptive immunity profiling. Nat Methods (2015) 12:380–381. doi: 10.1038/nmeth.3364

22. Shugay M, Bagaev D V., Turchaninova MA, Bolotin DA, Britanova O V., Putintseva E V., Pogorelyy M V., Nazarov VI, Zvyagin I V., Kirgizova VI, et al. VDJtools: Unifying Post-analysis of T Cell Receptor Repertoires. PLoS Comput Biol (2015) 11:1–16. doi: 10.1371/journal.pcbi.1004503

23. Jones E, Oliphant TE, Peterson P. SciPy: Open Source Scientific Tools for Python. (2001) Available at: http://www.scipy.org

24. Mahe F, Rognes T, Quince C, Vargas C De, Dunthorn M. Swarm: robust and fast clustering method for amplicon-based studies. PeerJ (2014) 2:1–13. doi: 10.7717/peerj.593

25. Huang H, Wang C, Rubelt F, Scriba TJ, Davis MM. Analyzing the Mycobacterium tuberculosis immune response by T-cell receptor clustering with GLIPH2 and genome-wide antigen screening. Nat Biotechnol (2020) doi: 10.1038/s41587-020-0505-4

26. Bastian M, Heymann S. Gephi : An Open Source Software for Exploring and Manipulating Networks. in

27. Sethna Z, Elhanati Y, Callan CG, Walczak AM, Mora T. OLGA: fast computation of generation probabilities of B- and T-cell receptor amino acid sequences and motifs. Bioinformatics (2018)1–8. doi: 10.1093/bioinformatics/btz035

28. Zheng GXY, Terry JM, Belgrader P, Ryvkin P, Bent ZW, Wilson R, Ziraldo SB, Wheeler TD, McDermott GP, Zhu J, et al. Massively parallel digital transcriptional profiling of single cells. Nat Commun (2017) 8: doi: 10.1038/ncomms14049

29. Butler A, Hoffman P, Smibert P, Papalexi E, Satija R. Integrating single-cell transcriptomic data across different conditions, technologies, and species. Nat Biotechnol (2018) 36:411–420. doi: 10.1038/nbt.4096

30. Mamedov MR, Scholzen A, Nair R V., Cumnock K, Kenkel JA, Oliveira JHM, Trujillo DL, Saligrama N, Zhang Y, Rubelt F, et al. A Macrophage Colony-Stimulating-Factor-Producing γδ T Cell Subset Prevents Malarial Parasitemic Recurrence. Immunity (2018) 48:350–363.e7. doi: 10.1016/j.immuni.2018.01.009

31. Venturi V, Kedzierska K, Turner SJ, Doherty PC, Davenport MP. Methods for comparing the diversity of samples of the T cell receptor repertoire. J Immunol Methods (2007) 321:182–195. doi: 10.1016/j.jim.2007.01.019

32. Britanova O V., Putintseva E V., Shugay M, Merzlyak EM, Turchaninova MA, Staroverov DB, Bolotin DA, Lukyanov S, Bogdanova EA, Mamedov IZ, et al. Age-Related Decrease in TCR Repertoire Diversity Measured with Deep and Normalized Sequence Profiling. J Immunol (2014) 192:2689–2698. doi: 10.4049/jimmunol.1302064

33. Izraelson M, Nakonechnaya TO, Moltedo B, Egorov ES, Kasatskaya SA, Putintseva E V., Mamedov IZ, Staroverov DB, Shemiakina II, Zakharova MY, et al. Comparative analysis of murine T-cell receptor repertoires. Immunology (2018) 153:133–144. doi: 10.1111/imm.12857

34. Madi A, Poran A, Shifrut E, Reich-zeliger S, Greenstein E, Zaretsky I, Arnon T, Laethem F Van, Singer A, Lu J, et al. T cell receptor repertoires of mice and humans are clustered in similarity networks around conserved public CDR3 sequences. Elife (2017) 3:1–17. doi: 10.7554/eLife.22057

35. Glanville J, Huang H, Nau A, Hatton O, Wagar LE, Rubelt F, Ji X, Han A, Krams SM, Pettus C, et al. Identifying specificity groups in the T cell receptor repertoire. Nature (2017) 547:94–98. doi: 10.1038/nature22976

36. Dash P, Fiore-Gartland AJ, Hertz T, Wang GC, Sharma S, Souquette A, Crawford JC, Clemens EB, Nguyen THO, Kedzierska K, et al. Quantifiable predictive features define epitope-specific T cell receptor repertoires. Nature (2017) 547:89–93. doi: 10.1038/nature22383

37. Covacu R, Philip H, Jaronen M, Douek DC, Efroni S, Quintana FJ, Covacu R, Philip H, Jaronen M, Almeida J, et al. System-wide Analysis of the T Cell Response. Cell Rep (2016) 14:2733–2744. doi: 10.1016/j.celrep.2016.02.056

38. Miles JJ, Bulek AM, Cole DK, Gostick E, Schauenburg AJA, Venturi V, Davenport MP, Tan MP, Burrows SR, Wooldridge L, et al. Genetic and Structural Basis for Selection of a Ubiquitous T Cell Receptor Deployed in Epstein-Barr Virus Infection. PLoS Pathog (2010) 6:1–15. doi: 10.1371/journal.ppat.1001198

39. Benati D, Galperin M, Lambotte O, Gras S, Lim A, Mukhopadhyay M, Nouël A, Campbell K, Lemercier B, Claireaux M, et al. Public T cell receptors confer high-avidity CD4 responses to HIV controllers. J Clin Invest (2016) 126:2093–2108. doi: 10.1172/JCI83792DS1

40. Madi A, Shifrut E, Reich-zeliger S, Gal H, Best K, Ndifon W, Chain B, Cohen IR, Friedman N. T-cell receptor repertoires share a restricted set of public and abundant CDR3 sequences that are associated with self-related immunity. Genome Res (2014) 24:1603–1612. doi: 10.1101/gr.170753.113.24

41. Venturi V, Kedzierska K, Price DA, Doherty PC, Douek DC, Turner SJ, Davenport MP. Sharing of T cell receptors in antigen-specific responses is driven by convergent recombination. PNAS (2006) 103:18691–18696. doi: 10.1073/pnas.0608907103

42. Pogorelyy M V., Minervina AA, Chudakov DM, Mamedov IZ, Lebedev YB, Mora T, Walczak AM. Method for identification of condition-associated public antigen receptor sequences. Elife (2018)1–13. doi: 10.7554/eLife.33050

43. Singhania A, Graham CM, Gabryšová L, Moreira-Teixeira L, Stavropoulos E, Pitt JM, Chakravarty P, Warnatsch A, Branchett WJ, Conejero L, et al. Transcriptional profiling unveils type I and II interferon networks in blood and tissues across diseases. Nat Commun (2019) 10:1–21. doi: 10.1038/s41467-019-10601-6

44. Elyahu Y, Hekselman I, Eizenberg-Magar I, Berner O, Strominger I, Schiller M, Mittal K, Nemirovsky A, Eremenko E, Vital A, et al. Aging promotes reorganization of the CD4 T cell landscape toward extreme regulatory and effector phenotypes. Sci Adv (2019) 5: doi: 10.1126/sciadv.aaw8330

45. Lönnberg T, Svensson V, James KR, Fernandez-Ruiz D, Sebina I, Montandon R, Soon MSF, Fogg LG, Nair AS, Liligeto UN, et al. Single-cell RNA-seq and computational analysis using temporal mixture modeling resolves TH1/TFH fate bifurcation in malaria. Sci Immunol (2017) 2:1–12. doi: 10.1126/sciimmunol.aal2192

46. Weiss GE, Traore B, Kayentao K, Ongoiba A, Doumbo S, Kone Y, Dia S, Guindo A, Traore A, Huang C, et al. The Plasmodium falciparum-Specific Human Memory B Cell Compartment Expands Gradually with Repeated Malaria Infections. PLoS Pathog (2010) 6: doi: 10.1371/journal.ppat.1000912

47. Emerson RO, DeWitt WS, Vignali M, Gravley J, Hu JK, Osborne EJ, Desmarais C, Klinger M, Carlson CS, Hansen JA, et al. Immunosequencing identifies signatures of cytomegalovirus exposure history and HLA-mediated effects on the T cell repertoire. Nat Genet (2017) 49:659–665. doi: 10.1038/ng.3822

48. Shugay M, Bagaev D V, Zvyagin I V, Vroomans RM, Crawford JC, Dolton G, Komech EA, Sycheva AL, Koneva AE, Egorov ES, et al. VDJdb?: a curated database of T-cell receptor sequences with known antigen specificity. Nucleic Acids Res (2018) 46:419–427. doi: 10.1093/nar/gkx760

49. Wolf K, Hether T, Gilchuk P, Ahn T, Joyce S, Dipaolo RJ, Wolf K, Hether T, Gilchuk P, Kumar A, et al. Identifying and Tracking Low-Frequency Virus-Specific TCR Clonotypes Using High-Throughput Sequencing. Cell Rep (2018) 25:2369–2378.e4. doi: 10.1016/j.celrep.2018.11.009

50. Su Z, Stevenson MM. Central role of endogenous gamma interferon in protective immunity against blood-stage Plasmodium chabaudi AS infection. Infect Immun (2000) 68:4399–4406. doi: 10.1128/IAI.68.8.4399-4406.2000

51. Soon MSF, Haque A. Recent Insights into CD4 + Th Cell Differentiation in Malaria. J Immunol (2018) 200:1965–1975. doi: 10.4049/jimmunol.1701316

52. Pérez-Mazliah D, Ng DHL, Freitas do Rosário AP, McLaughlin S, Mastelic-Gavillet B, Sodenkamp J, Kushinga G, Langhorne J. Disruption of IL-21 Signaling Affects T Cell-B Cell Interactions and Abrogates Protective Humoral Immunity to Malaria. PLoS Pathog (2015) 11:1–24. doi: 10.1371/journal.ppat.1004715

53. Pérez-Mazliah D, Nguyen MP, Hosking C, McLaughlin S, Lewis MD, Tumwine I, Levy P, Langhorne J. Follicular Helper T Cells are Essential for the Elimination of Plasmodium Infection. EBioMedicine (2017) 24:216–230. doi: 10.1016/j.ebiom.2017.08.030

54. Tubo NJ, Pagán AJ, Taylor JJ, Nelson RW, Linehan JL, Ertelt JM, Huseby ES, Way SS, Jenkins MK. Single naive CD4+ T cells from a diverse repertoire produce different effector cell types during infection. Cell (2013) 153:785–796. doi: 10.1016/j.cell.2013.04.007

55. Schattgen SA, Guion K, Crawford JC, Souquette A, Barrio AM, Stubbington MJT, Thomas PG, Bradley P. Linking T cell receptor sequence to transcriptional profiles with clonotype neighbor graph analysis (CoNGA). bioRxiv (2020)2020.06.04.134536. doi: 10.1101/2020.06.04.134536

56. Khosravi-Maharlooei M, Obradovic A, Misra A, Motwani K, Holzl M, Seay HR, DeWolf S, Nauman G, Danzl N, Li H, et al. Crossreactive public TCR sequences undergo positive selection in the human thymic repertoire. J Clin Invest (2019) 130: doi: 10.1172/JCI124358

57. Miles JJ, Douek DC, Price DA. Bias in the α? T-cell repertoire: Implications for disease pathogenesis and vaccination. Immunol Cell Biol (2011) 89:375–387. doi: 10.1038/icb.2010.139

58. Gasper DJ, Tejera MM, Suresh M. CD4 T-cell memory generation and maintenance. Crit Rev Immunol (2014) 34:121–46. doi: 10.1615/critrevimmunol.2014010373

59. Tickotsky N, Sagiv T, Prilusky J, Shifrut E, Friedman N. McPAS-TCR: A manually curated catalogue of pathology-associated T cell receptor sequences. Bioinformatics (2017) 33:2924–2929. doi: 10.1093/bioinformatics/btx286

60. Hill DL, Pierson W, Bolland DJ, Mkindi C, Carr EJ, Wang J, Houard S, Wingett SW, Audran R, Wallin EF, et al. The adjuvant GLA-SE promotes human Tfh cell expansion and emergence of public TCRβ clonotypes. J Exp Med (2019)jem.20190301. doi: 10.1084/jem.20190301

