## Supplementary Figures for "A Conserved TCRβ signature dominates a highly polyclonal T-cell expansion during the acute phase of a murine malaria infection"

### Supplementary Figure 1

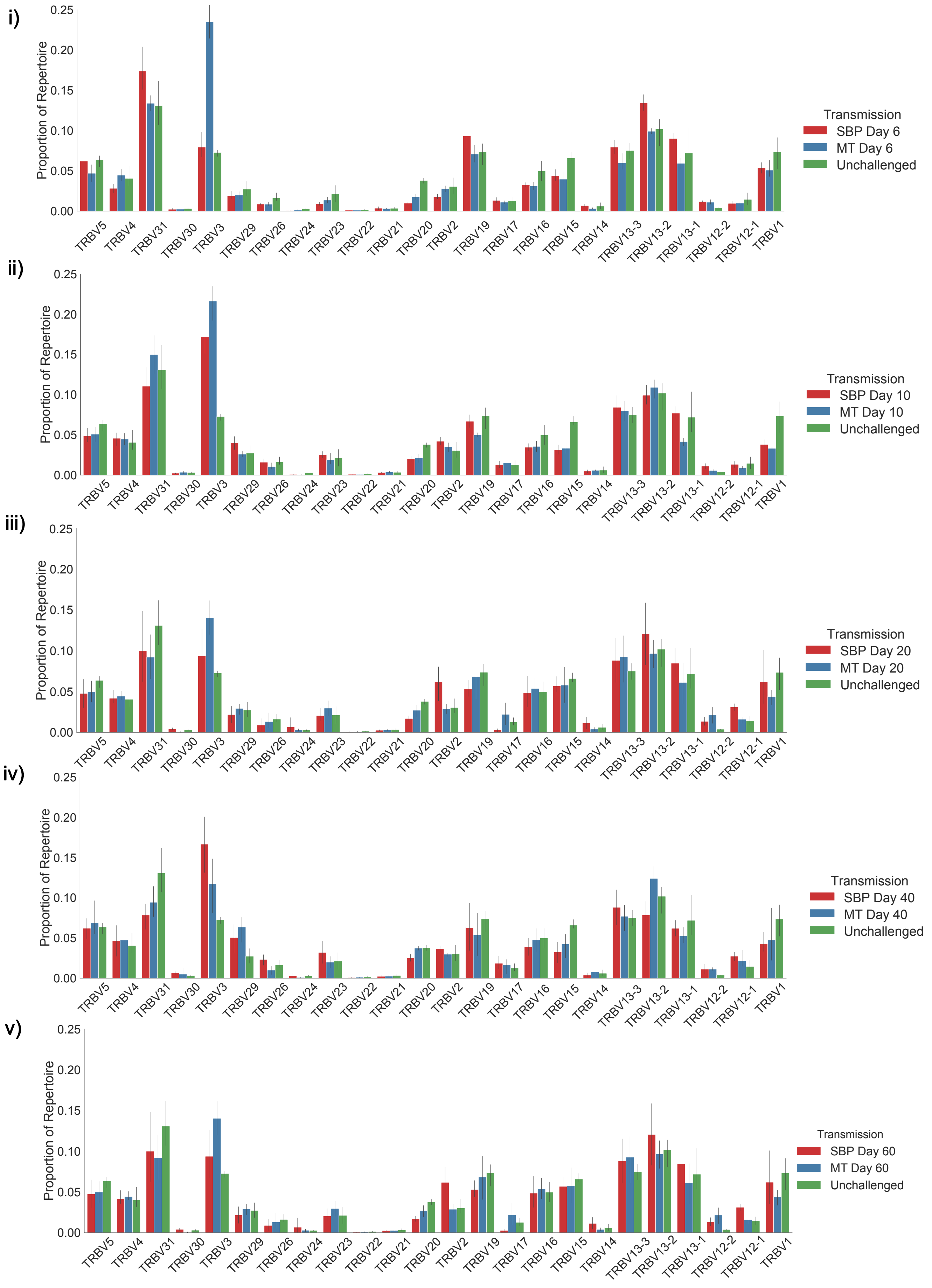


**Supplementary Figure 1:** Proportion of TRBV-gene usage in unchallenged $T_N$ repertoires (green), and $T_E$ repertoires of mice infected with SBP parasites (red) or recently MT parasites (blue) at days 6 (i), 10 (ii), 20 (iii), 40 (iv) and 60 (v) days post-infection.

Supplementary Figure 2

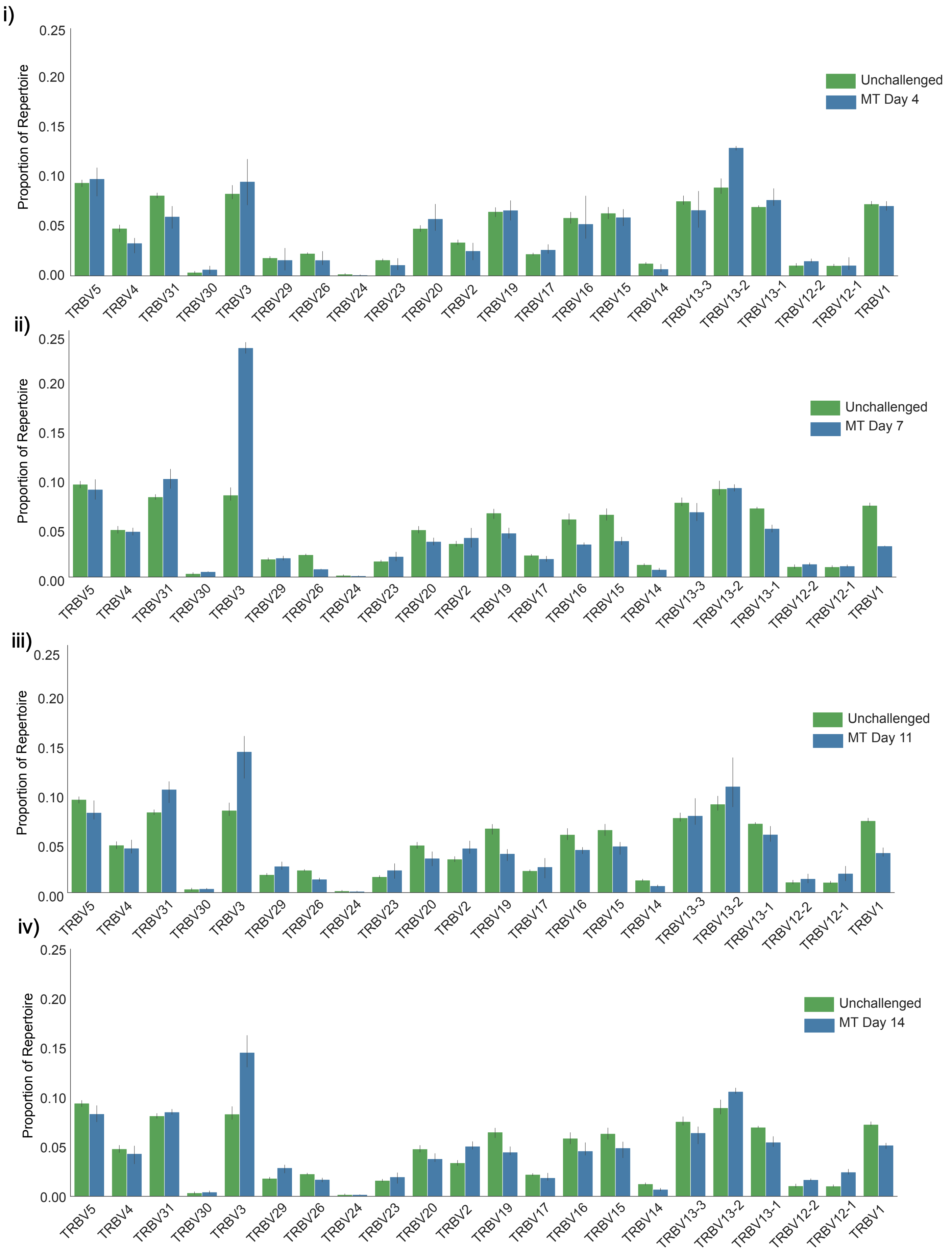


**Supplementary Figure 2:** Proportion of TRBV-gene usage in unchallenged $T_N$ repertoires (green) and $T_E$ repertoires of mice infected with recently MT parasites (blue) in a second independent experiment at days 4(i), 7 (ii), 11 (iii) and 14 (iv) post-infection.

Supplementary Figure 3

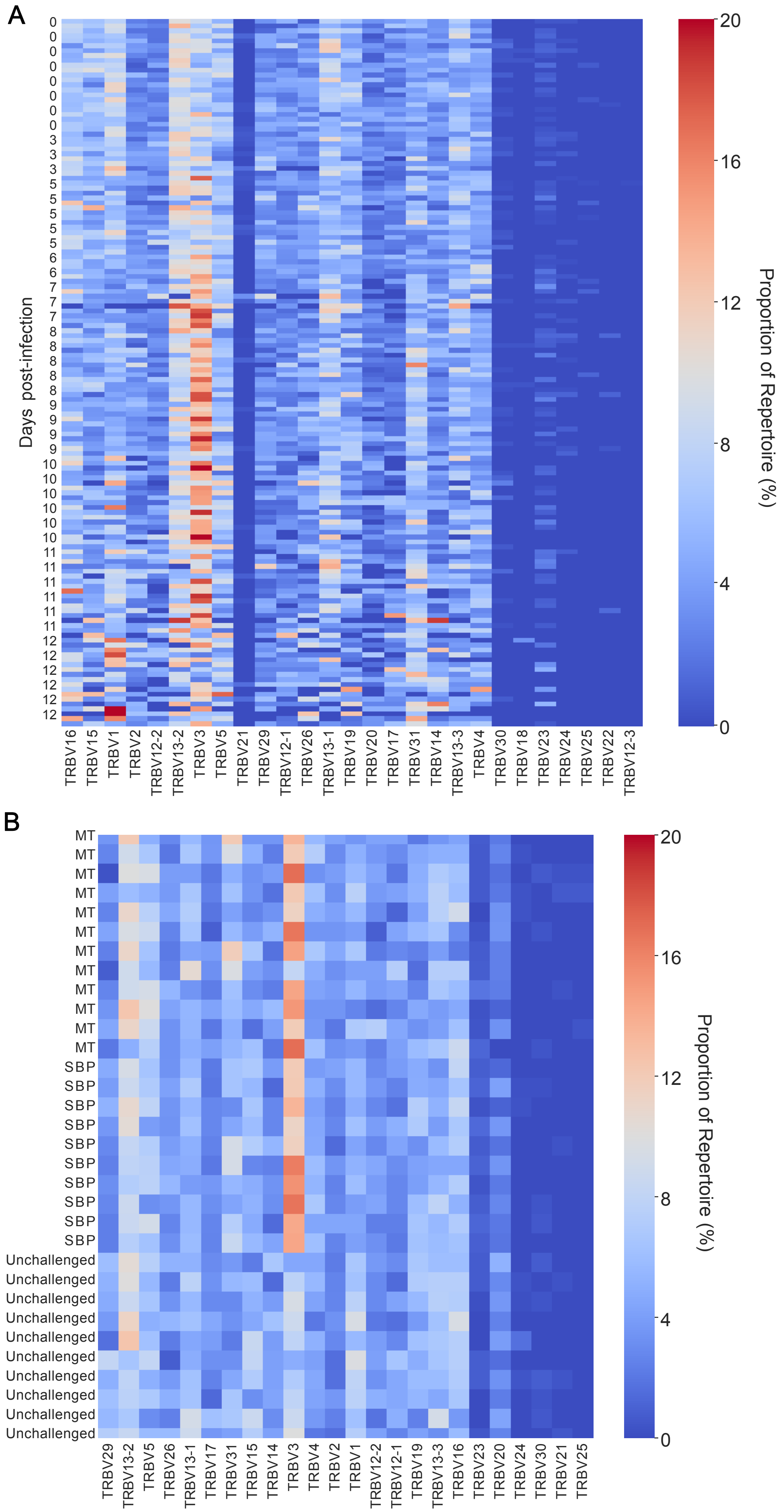


**Supplementary Figure 3:** Heatmaps show proportion of TRBV-gene usage in unchallenged and challenged mice unsorted splenic TCRβ repertoires from C57BL/6 mice infected with bloodstage A) *P. chabaudi (AS)* and B) *P. chabaudi (CB)*. TCRβ repertoires were reconstructed from publicly available RNA-seq data sets. Each column represents a unique TRBV gene, and each row is an individual replicate mouse. No time-point data is available for (B).

Supplementary Figure 4

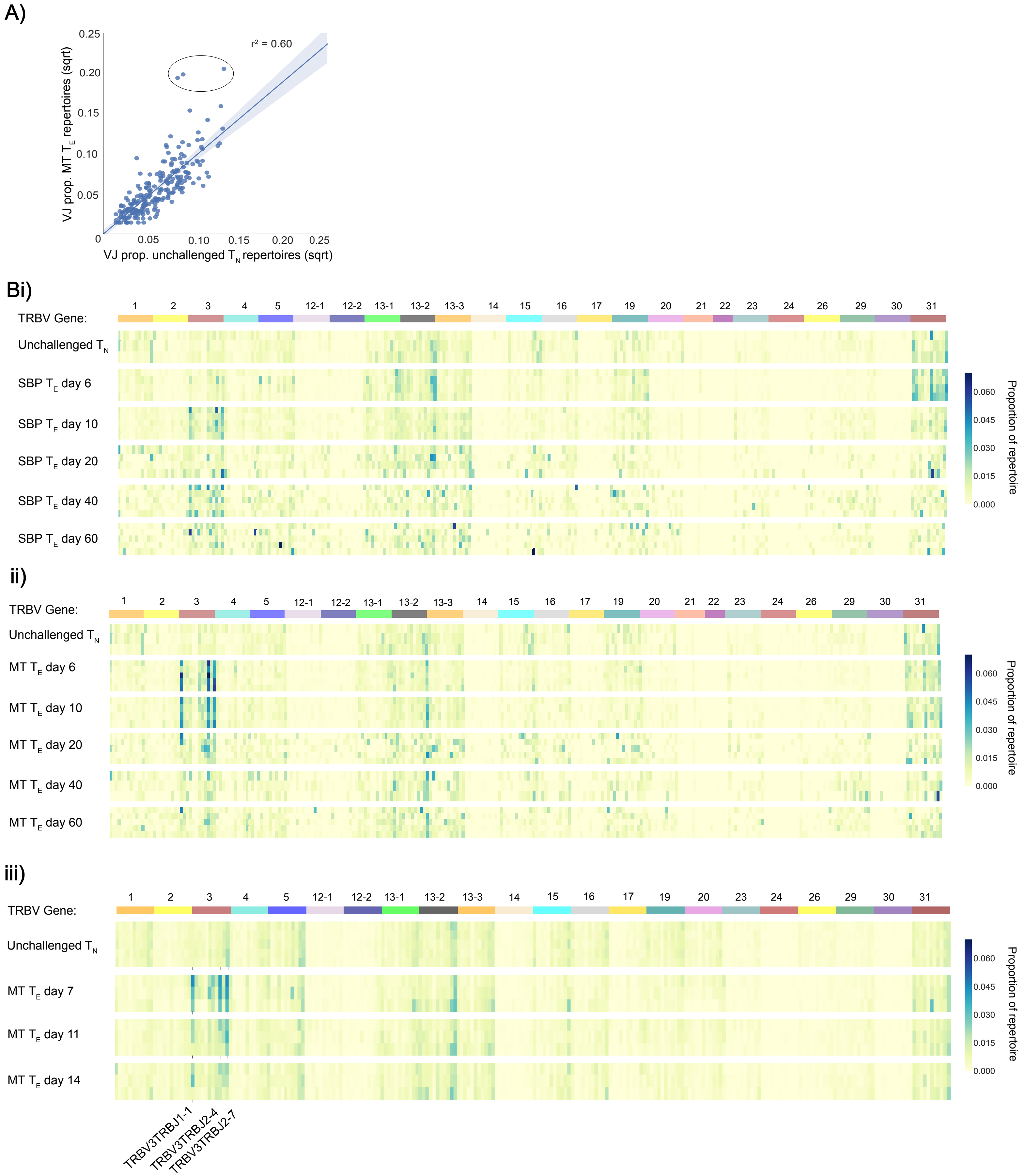


**Supplementary Figure 4:** Mean proportion of each V/J allelic combination in unchallenged $T_N$ repertoires versus challenged $T_E$ repertoires at days 7 (i) post-infection for mice infected with recently MT parasites in a second independent experiment. B) Heatmap depicts proportion of each V/J allelic combination (columns) for individual replicate mouse (rows) for (i) unchallenged $T_N$ repertoires and acute $T_E$ SBP repertoires and (ii) unchallenged $T_N$ repertoires and acute $T_E$ MT repertoires and (iii) unchallenged $T_N$ repertoires and acute $T_E$ MT repertoires from a second independent experiment.

Supplementary Figure 5

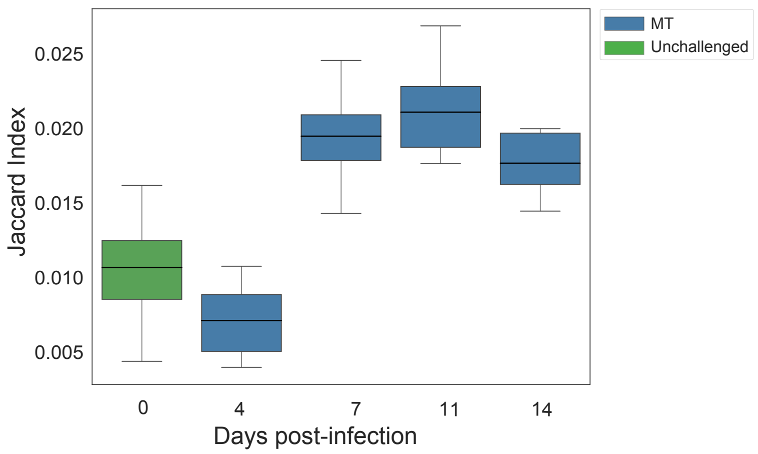


**Supplementary Figure 5:** Jaccard similarity index of unchallenged $T_N$ repertoires (green) and $T_E$ repertoires of mice infected with recently MT parasites (blue) in a second independent experiment at days 4, 7, 11 and 14 post-infection. Data was normalised by down-sampling to 5000 UMI.

**Supplementary Table 1:** Summary results for GLIPH2 analysis. Filter criteria used: CDR3 sequence present in at least 75% of replicate group; vb_score<=0.05; length_score <=0.05; expansion_score <=0.05; Fisher_score <=0.001

| Infection type | Time point | No. of clusters meeting criteria | Cluster_ID | Pattern | Final_score | |
|---|---|---|---|---|---|---|
| MT | D6 | 5 | 14 | S%SQNT | 8.50E-14 | OTU1 |
| | | | 18 | SPT%NTE | 3.10E-13 | |
| | | | 10 | SLS%NTE | 4.60E-13 | |
| | | | 4 | SLTQ | 4.60E-12 | |
| | | | 7 | SL%QGAE | 8.50E-12 | |
| MT | D10 | 4 | 21 | S%SQNT | 7.70E-14 | OTU1 |
| | | | 32 | S%GQYE | 4.40E-13 | OTU2 |
| | | | 5 | SLS%NTE | 1.70E-12 | |
| | | | 23 | SL%QNTE | 6.10E-12 | |
| MT | D20 | 0 | | | | |
| MT | D40 | 0 | | | | |
| MT | D60 | 0 | | | | |
| SBP | D6 | 0 | | | | |
| SBP | D10 | 1 | 16 | S%GTTSAET | 1.10E-09 | |
| SBP | D20 | 0 | | | | |
| SBP | D40 | 0 | | | | |
| SBP | D60 | 0 | | | | |

Supplementary Figure 6

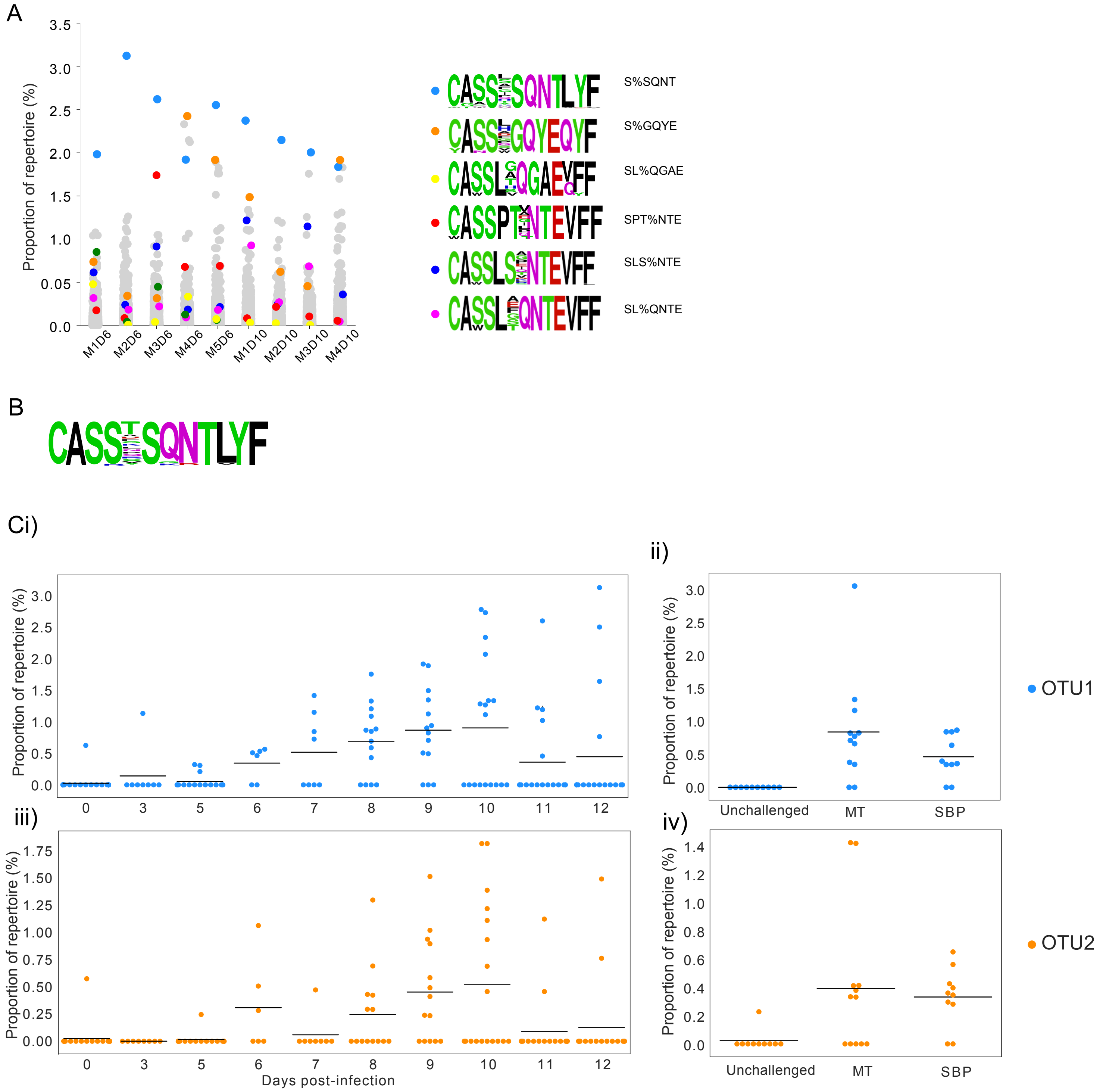


**Supplementary Figure 6:** Strip plots indicate proportion of individual repertoires taken up by clusters identified as significantly enriched using GLIPH2 (results in Supplementary Table 1), with representative sequence logos for each cluster. B) Representative amino acid sequence logo of the most dominant cluster detected (OTU1) from a second independent experiment. C) Proportion of CDR3s in cluster OTU1 (blue) and OTU2 (orange) found in unsorted splenic TCRβ repertoires reconstructed from publicly available RNA-seq data for i) and iii) *P. chabaudi (AS),* and for ii) and iv) *P. chabaudi (CB).* No time point data is available for the *P.chabaudi* (CB) dataset. Black horizontal line indicates mean, and each point represents an individual mouse.

Supplementary Figure 7

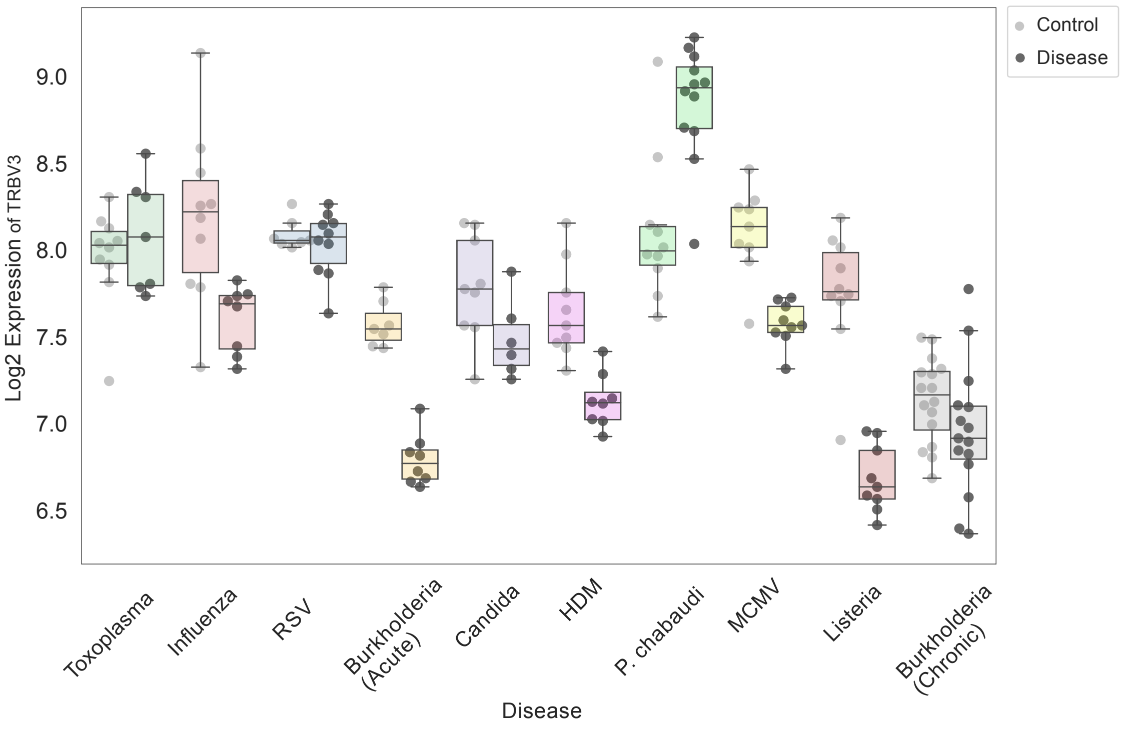


**Supplementary Figure 7:** Log2 expression values (normalised RNA-seq counts) of TRBV3 from publicly available whole blood RNA-seq data from C57Bl/6 mice, taken at peak of murine response for each disease. Data is paired as unchallenged controls (light grey circles) and challenged (black circles).

Supplementary Figure 8

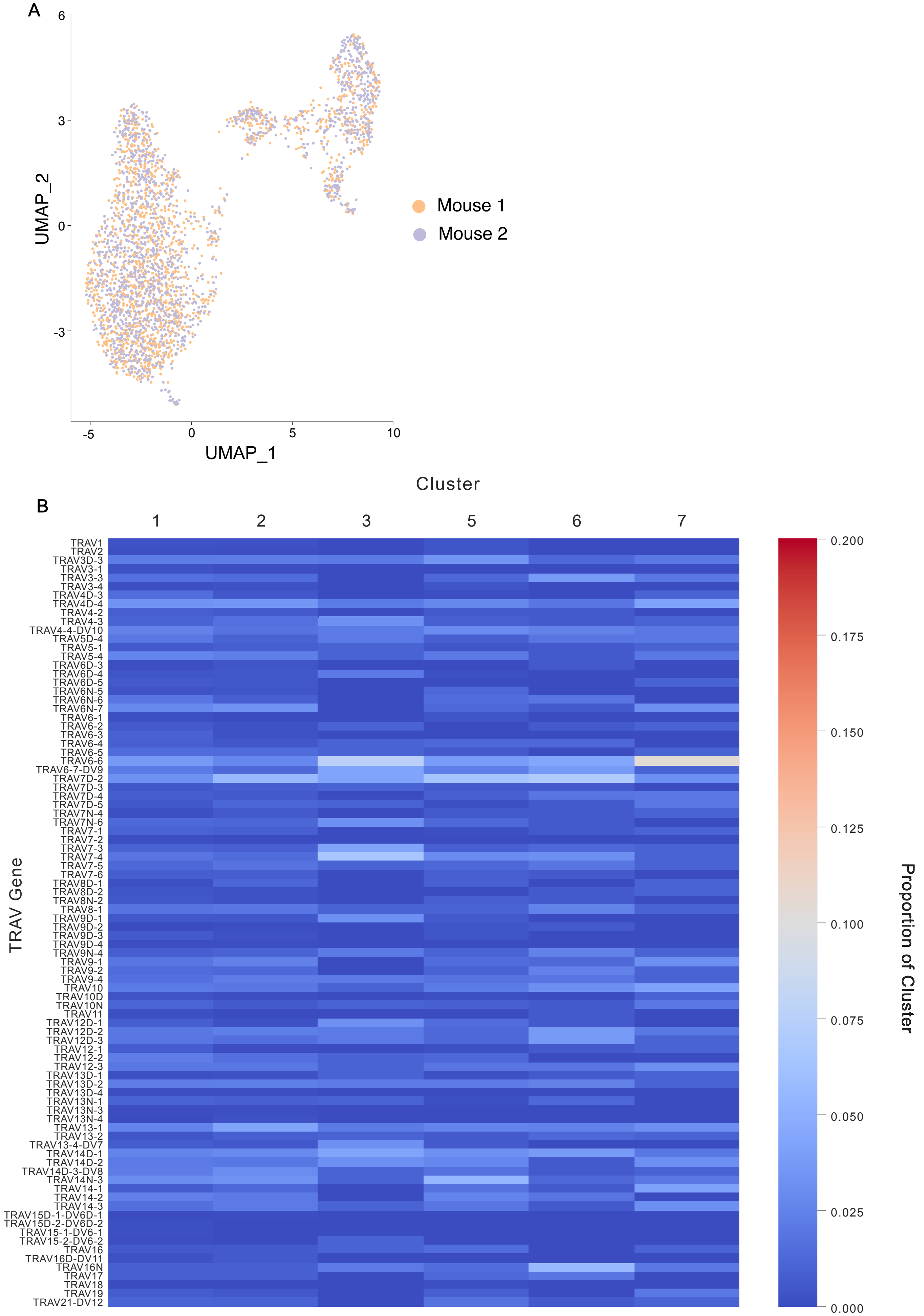


**Supplementary Figure 8:** A) Two-dimensional UMAP visualization of CD4⁺ splenocytes from 2 challenged mice, at day 7 post-infection, as per Seurat cluster assignment, with cells coloured by individual; cell positions are as per UMAP in Figure 8A. B) Heatmap displays proportion of TRAV gene per cluster TCR repertoire.

### Supplementary Figure 9

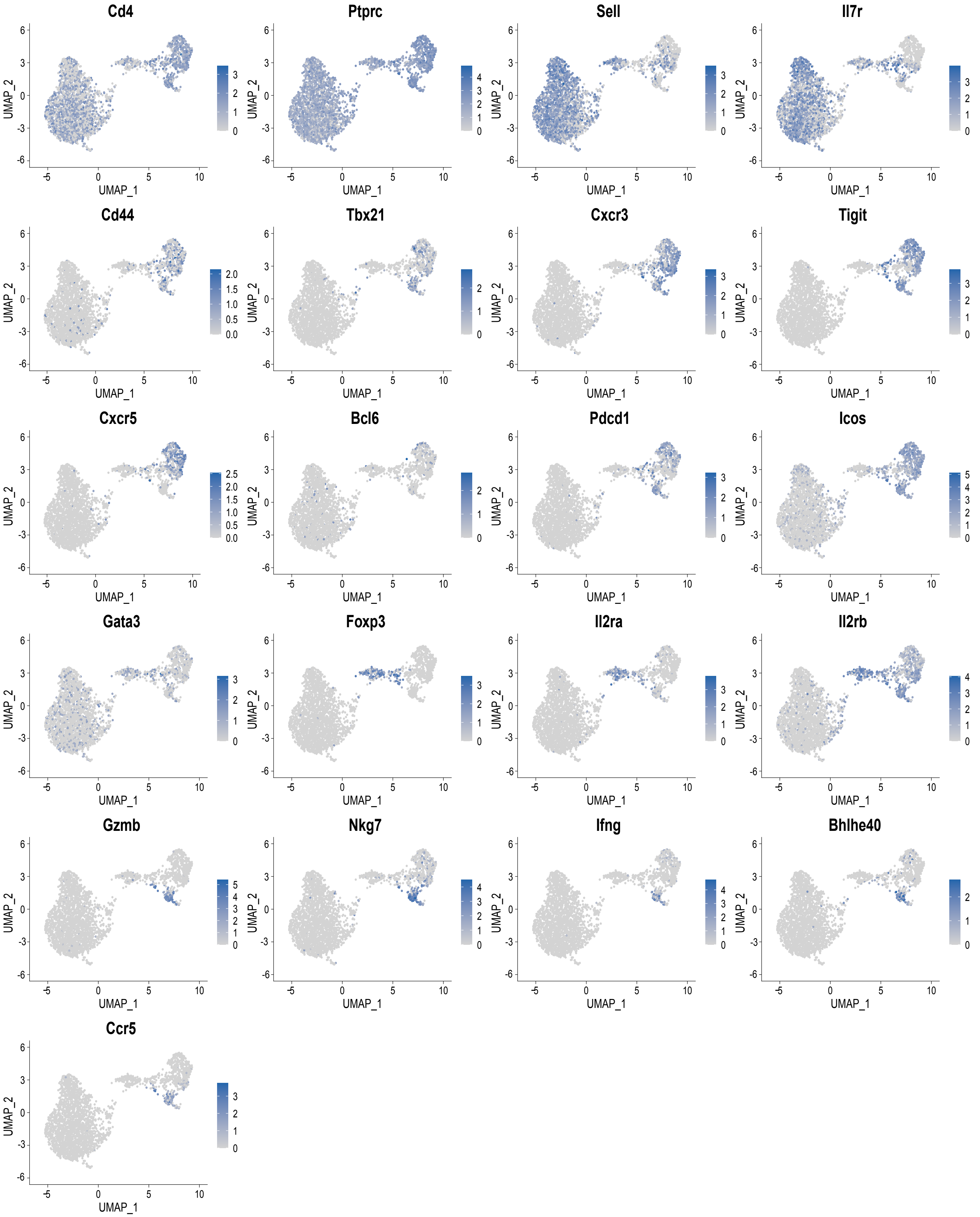


**Supplementary Figure 9:** Expression of T-cell marker genes of interest, from CD4⁺ splenocytes of 2 challenged mice at day 7 post *P. chabaudi* infection; cell positions are as per UMAP in Figure 8A.

Supplementary Figure 10

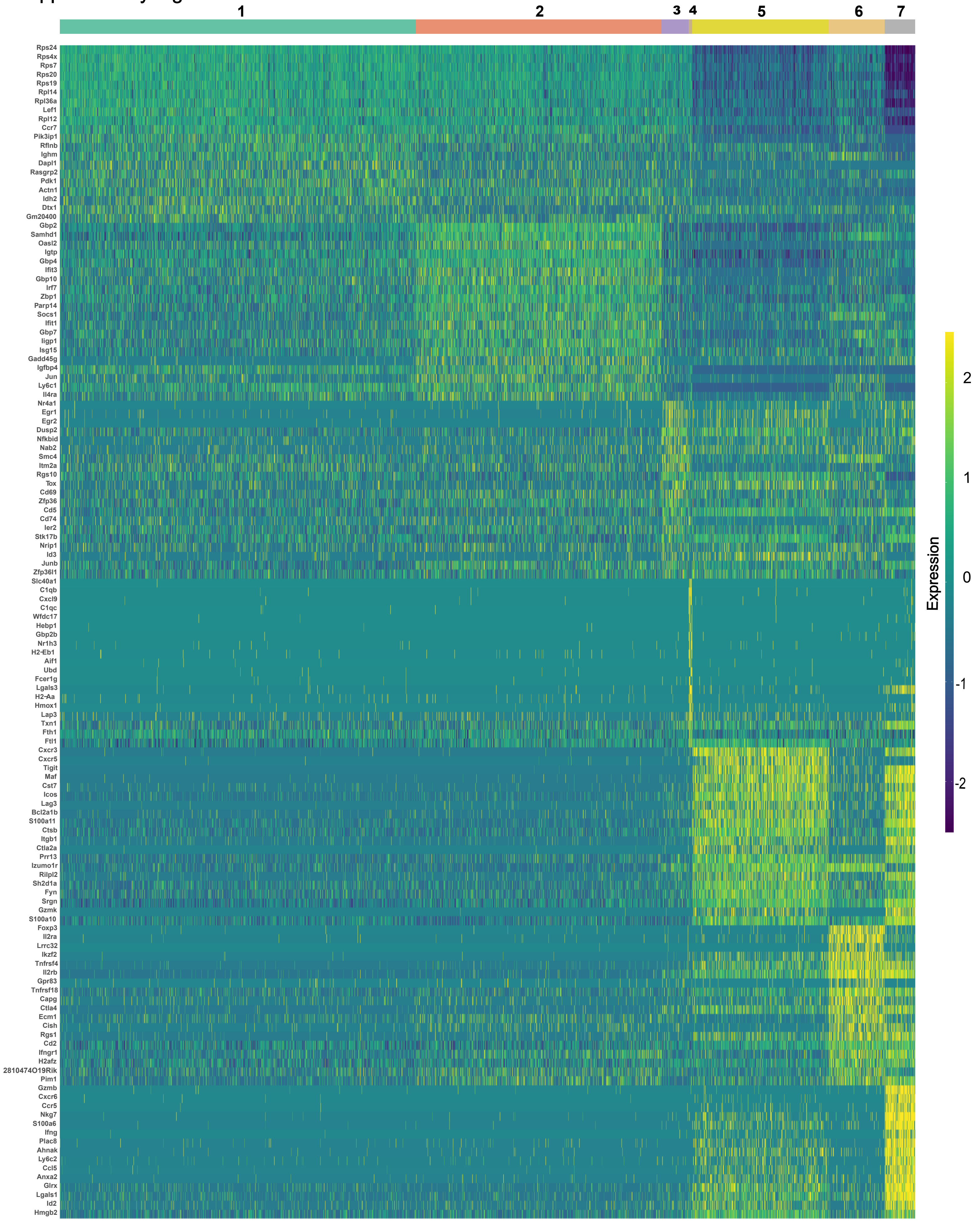


**Supplementary Figure 10:** D) Heatmap depicts the top 20 differentially expressed genes and their expression values between all 7 distinct clusters, of CD4+ splenocytes of 2 challenged mice at day 7 post *P. chabaudi* infection.

Supplementary Figure 11

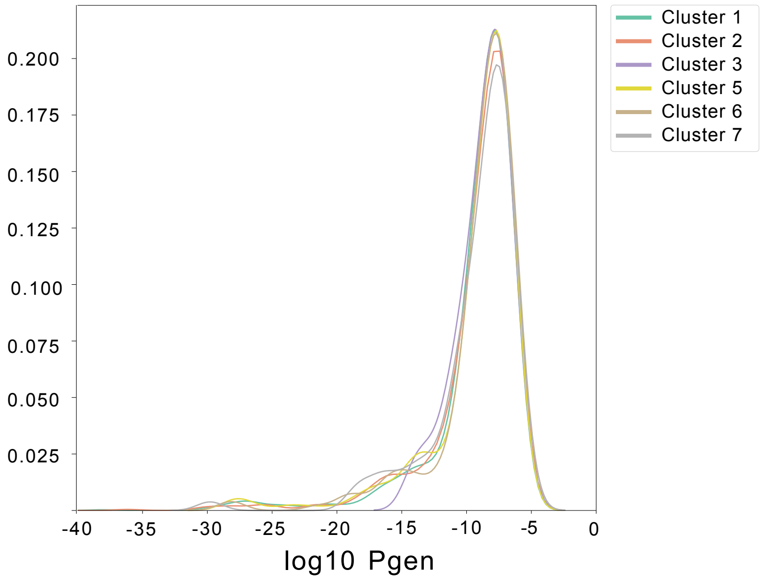


**Supplementary Figure 11:** KDE plot of the probability of generation (log10) of TCRβ CDR3 nucleotide sequences per cluster, of CD4⁺ splenocytes of 2 challenged mice at day 7 post *P. chabaudi* infection.
